# Biophysical resetting of specialization enables metabolic network diversification

**DOI:** 10.64898/2026.10.14.759191

**Authors:** Alexander V. Badyaev, Carmen Sánchez Moreno, Branden Lau, Erin S. Morrison

## Abstract

Organisms must integrate external inputs whose availability they do not control in internal functions that require precision. Birds exemplify this challenge: they cannot synthesize carotenoids, yet closely integrate diet-derived carotenoids into vision, pigmentation and yolk provisioning, raising the question of how unpredictability of external supply coevolves with precision of internal use. Here we analyze core carotenogenesis proteins across 172 bird species and show that the resolution is a biophysical bow-tie in which ordered, self-attracting absorption and enzymatic complexes are linked by disordered, phase-separating and self-avoiding carriers. This architecture interrupts the propagation of upstream specialization without compromising downstream specificity: nonspecific connectors allow diverse inputs to access and recombine among functional modules. Across species, the biophysical properties of these connectors, but not of conserved absorption or enzymatic complexes, drive network diversification through downstream metabolic branching rather than the accumulation of dedicated input-specific pathways. Decoupling the upstream origin of inputs from their downstream routing by the biophysical bow-tie architecture accounts for three defining features of avian carotenogenesis: physiological sensitivity, openness to novel inputs, and the internalization of metabolic control. Assembly of non-specific carriers thus produce specific outcomes, sustaining the connectivity and remarkable evolutionary diversification of a metabolic network built entirely on an unpredictable external resource.

**Significance:** How can organisms evolve precise functions that depend on unpredictable external resources? Dedicated connections between inputs and outputs would propagate environmental contingency, hindering evolution of internal functions. We identify an alternative in avian carotenoid metabolism: specialized functional complexes are linked by a biophysically distinct carrier layer that interrupts the propagation of upstream specialization without eliminating downstream specificity. Across species, the properties of this resetting layer predict diversification through branching among conserved metabolic complexes rather than the accumulation of dedicated input– output connections. The ability to interrupt the propagation of specialization may therefore be as important for evolutionary diversification as the ability to generate specialization itself.

## Main

Gains and losses of external dependencies drive organismal evolution (1–4), requiring regulatory systems able to recognize and control external inputs whose availability is unpredictable. Organisms thus face the trade-off between internal specialization and external unpredictability of resources on which they depend, raising a question of how specialized uptake of external inputs coevolves with precision of their internal utilization. A dedicated, input-specific intermediate regulator would preserve specificity, but propagate fluctuations of external inputs, thus hindering evolution of their internal integration (Fig. 1a). Decoupling upstream origin from downstream utilization by non-specific intermediates would buffer supply variability but requires additional mechanisms to restore downstream specificity of inputs (5, 6). A resolution is suggested by recent empirical work on emerging specificity in regulatory mechanisms (7–9), where a combination of phase-separation propensity and binding self-avoidance allow carrier proteins to transport diverse chemical compounds while preserving their functional identity. In such a system, cargoes remain segregated (e.g., by physicochemical recognition) in discrete non-coalescing vehicles and independently routable even though the carriers themselves are chemically indiscriminate (10–13). Specificity of downstream delivery in such a system could arise from constraints imposed by assemblies of non-specific channels of transmission themselves (9, 14) (Fig. 1c). In essence, these processes *decontextualize* external inputs, capitalizing on changes in modality of their transmission (see also (15)).

**Fig. 1.**
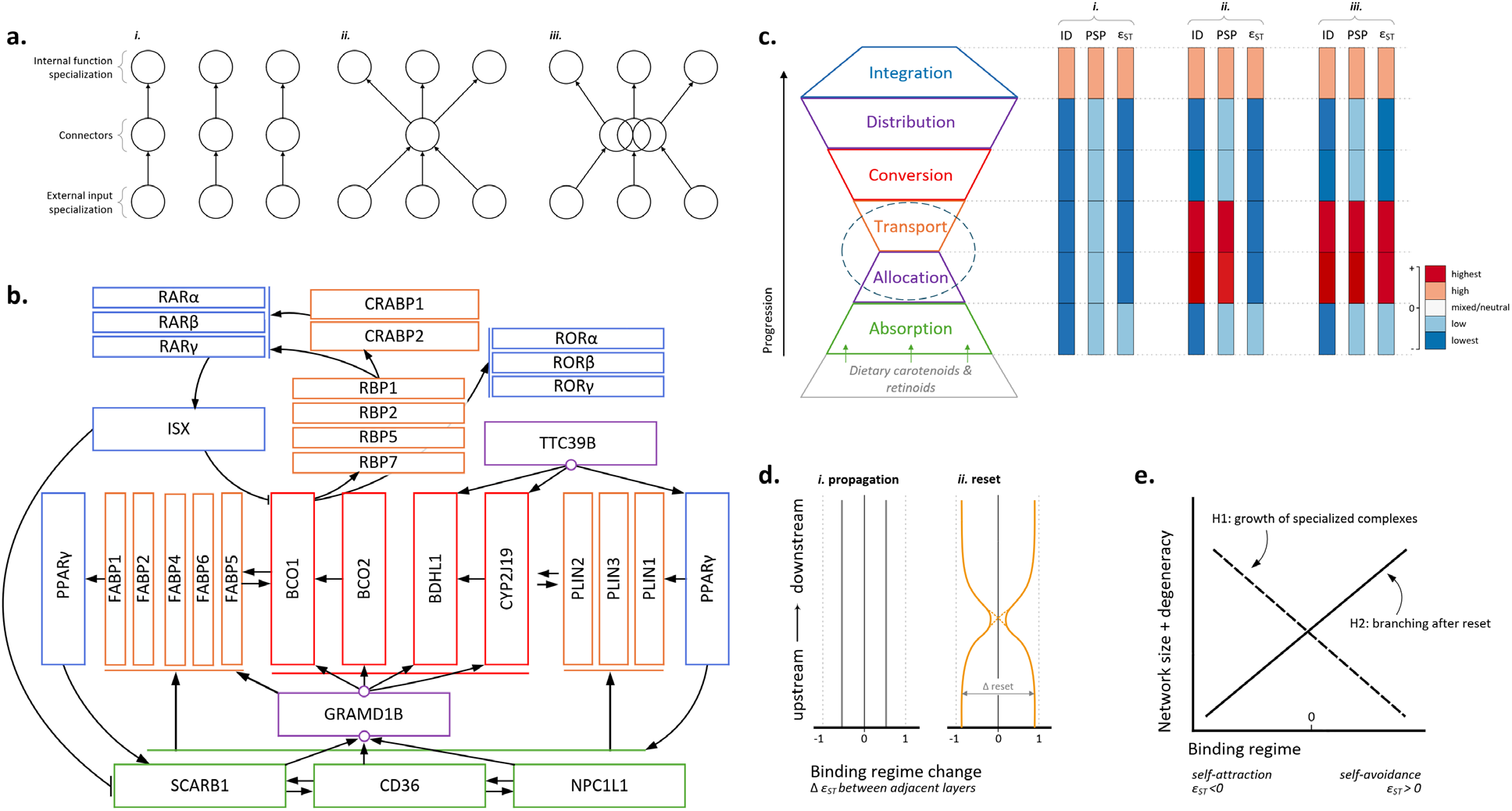
Converting diversity and contingency of external inputs into internal precision. **(a)** Three scenarios: *i.* dedicated input-specific connectors transmit volatility of external inputs; *ii.* degenerate connectors buffer inputs’ lability, but compromise their specificity, *iii.* decoupling upstream origin from downstream utilization in external inputs while preserving identity of inputs resolves the trade-off. **(b)** Core avian carotenogenesis proteins (Supplementary Text S1, Supplementary Table S1) grouped into six functional groups: 1) uptake of dietary precursors (**absorption**, green); 2) pre-conversion (**allocation**, GRAMD1B) and 3) post-conversion (**distribution**, TTC39B) partitioning of products (violet); 4) cleavage and chemical **conversion** (red); 5) carriers (orange) that include within-cell (FABPs, RBPs, and CRABPs) and longer distance (PLINs) **transport and storage** proteins, and 6) **integration** (blue) – transcriptional regulators and cofactors. PPARγ is drawn on both sides of the network to show its interactions with FABP1, PLIN1, and TTC39B. **(c)** Distinct conformational stability and binding propensity of proteins in these groups can form a biophysical bow-tie architecture of this network, where intrinsic disorder (ID) and self-avoiding binding of proteins in **allocation**, **transport** and **integration** complexes can compartmentalize specialized **absorption** and **conversion** modules, while transmitting their products. Shown are predicted ID, phase-separation propensity (PSP) and homotypic binding propensity (ε_ST_ < 0 indicate self-attraction, ε_ST_ > 0 – self-avoidance) corresponding to *i-iii* scenarios in (a). Dashed ellipse outlines the waist layers of the proposed bow-tie architecture. **(d)** Propensity of specialized functional complexes to propagate (grey lines) or compartmentalize (gold lines) along the network pathways is reflected by the change in sign and magnitude of binding regime between adjacent layers: *i.* propagation produces consistent ε_ST_ regime, whereas *ii.* resetting compartmentalizes this specialization, representing a waist of the bow-tie architecture. **(e)** Evolutionary diversification of carotenoid networks can be driven by accumulation or expansion of specialized functional complexes (e.g., of absorption or conversion; dashed line; H1) or by branching following interruption of specialization propagation (solid line, H2).

In addition to the decontextualization, two additional requirements accommodate dynamic fluctuations of diverse external inputs (16). The first is the necessity to buffer against their temporal fluctuations, which favors evolution of storage capacity: the carriers could vary in their availability or the capacity to sequester and store their cargoes (17, 18). The second is the necessity to dynamically allocate fluctuating inputs to the carriers without interfering with upstream acquisition machinery that might be tuned to particular inputs. The resulting architecture would decouple the two specialized layers – acquisition of external inputs and internal functions – and thus facilitate evolution of precision in each – while also enabling changeability and robustness of the system by flexibly rerouting substrates between them by non-specialized carriers (Fig. 1a, *iii*)(16, 19).

Capitalizing on these insights, we examine whether such organization underpins evolution of the close control and precision of essential carotenoid-based functions in animals that cannot synthesize carotenoids *de novo* – a particularly tractable system that requires dynamic internalization of strictly external inputs (20–23). Despite wide environmental fluctuations in dietary carotenoids, birds closely integrate diverse diet-derived carotenoids into a multitude of essential and precise functions – from vision to pigmentation and immunity – as well as upregulate circulation of specific carotenoids during seasonal molt and oogenesis and channel them in a tissue-, sex-, and season-specific manner (24–31). Because the external supply fluctuates widely, this integration and its internal control depend on biochemical degeneracy – the ability to produce the same internal compound from different dietary precursors – which stabilizes internal function despite unstable inputs. In birds, this degeneracy sustains evolutionary subsampling and recombination of conserved network motifs, underpins close coevolution of external carotenoids and feather structure, and keeps the network poised for ecological exploration of novel carotenoids (32–35). Yet the regulatory basis of this degeneracy – where on the network it originates and what mechanism sustains it – is unknown. Degeneracy can be an outcome of accumulating multiple specialized biochemical paths linking external inputs and internal functions. Alternatively, it can be produced by resetting links, such as non-specialized carriers, among specialized modules (Fig. 1) (36).

To distinguish these scenarios, we first propose an architectural framework for the avian carotenogenesis network by delineating two specialized protein complexes (Fig. 1b; Supplementary Table S1, Supplementary Text S1): a membrane absorption layer needed to recognize, select and import an array of external carotenoids and an enzymatic conversion layer that generates a range of tissue-, sex-, and season-specific products. Ecological fluctuations of external carotenoids should prevent evolution of dedicated carrier linkages between these specialized layers (Fig. 1 a, *i*) (37). Instead, these specialized layers can be linked by an intermediate layer of carriers that can bind carotenoids non-specifically and transfer them to any downstream module (the bow-tie organization hypothesis, Fig. 1c). This architecture would account for two key features of avian carotenoid networks: degeneracy – interchangeable routes from inputs to outputs, which sustains persistence despite dependency on unpredictable external inputs, and continuity, which keeps a network connected as its motifs are subsampled and recombined over development and evolution (33).

Here we propose that the two lipid-transfer complexes – the transport/storage proteins and the allocation proteins – comprise this intermediate resetting layer (Fig. 1c, Supplementary Text S1). Testing this hypothesis requires showing that this layer’s biophysical properties are distinct from those of specialized layers and conducive to the proposed resetting hub properties (Fig. 1c, d) with distinct consequences for network evolution (Fig. 1e). We specifically focus on three sequence-derived biophysical properties of proteins particularly relevant for enzymatic networks (38): intrinsic disorder (ID), droplet-forming phase-separation propensity (PSP), and the sign and strength of self-referential (homotypic) binding (ε_ST_ < 0 denotes self-attraction with propensity to build dedicated same-partner complexes, whereas ε_ST_ > 0 denotes self-avoidance, which prevents these complexes) (9, 39).

The combination of layer-specific protein profiles allows us to distinguish three scenarios for the biophysical organization of the network (Fig. 1a, c). All scenarios predict that proteins comprising the specialized layers – e.g., absorption, conversion, post-conversion distribution – should be ordered, weakly phase-separating, and self-attracting, forming cooperative functional complexes whose specialization propagates along the biochemical axis (Fig. 1d, *i*), whereas proteins comprising the integration layer should be disordered, phase-separating, and self-avoiding, forming the transient transcription co-activator and co-repressor condensates. The distinction between the three scenarios is the predicted properties of transport and allocation proteins linking the specialized modules (hereafter “the waist” of the bow-tie architecture – dashed ellipse in Fig. 1c). Under the dedicated connector scenario (Fig. 1a, *i*), these proteins should be ordered and self-attracting, matching the specialized upstream and downstream layers they link, allowing propagation of specialization (Fig. 1c, d; *i*). In contrast, the bow-tie organization (Fig. 1a, scenarios *ii-iii*) predicts a reset at the waist (Fig. 1d, *ii*), such that transport and allocation proteins should be disordered and phase-separating (Fig. 1c). The decoupling of origin identity and downstream routing scenario (Fig. 1a, *iii*) specifically predicts the highest self-avoidance in these proteins compared to all others in the network (Fig. 1c, *iii*). In such an organization, self-avoidance reduces the persistence of dedicated same-partner assemblies, while heterotypic interactions and PSP-induced compartmentalization determine where and when carriers transfer or sequester their cargo.

These scenarios predict distinct patterns of evolution of avian carotenoid networks (Fig. 1e). Under the growth of specialized complexes model (H1), the self-attracting linker proteins build dedicated complexes and propagate specialization, accumulating specialized input-output modules (Fig. 1d, *i*). In contrast, the self-avoiding linker proteins reset the propagation of specialization between the input and output layers (H2, Fig. 1d, *ii*). Under the former scenario, network diversification is driven by gain and loss of specialized input-output modules and the capacity of a network to retain these modules (40). Under the latter scenario, network diversification is driven by recombination of pathways screened off their origin identity at the reset step (“branching after reset” scenario, Fig. 1e); the resulting degeneracy itself maintains network cohesion.

We test these predictions across the carotenogenesis networks of 172 bird species, quantifying the biophysical profile of 31 core proteins, tracing how specificity propagates and is reset across functional layers, and testing whether the properties of the waist hub scale with the size, connectivity and degeneracy of the carotenoid networks they sustain.

## Results

### Biophysical profiles reveal a functional bow-tie in the avian carotenogenesis

Across species, the three measures of disorder and binding attraction reveal the biophysical bow-tie organization (Figs. 1c, 2): ordered and self-attracting specialized complexes of absorption and conversion linked by disordered, phase-separating and self-avoiding carriers of their products. The carotenoid-processing enzymes in the conversion layer are ordered (with disorder confined to the membrane-bound segments of CYP2J19 and BDH1L, Supplementary Data S1 and S3) and weakly phase-separating; the two enzymes that build the dedicated ketolation and disposal machinery, CYP2J19 and BCO2, show strong self-attraction, reflecting their specialized membrane-associated clusters (Fig. 2a-c; Supplementary Text S1). Consistent with the functional separation of the oxidative and cleaving routes (Fig. 1b; Supplementary Text S1), BCO1 and BDH1L are not predicted to cluster (Fig. 2c). The dietary absorption layer (SCARB1, CD36, and NPC1L1) is similarly highly structured and weakly phase-separating (only NPC1L1 shows moderate phase separation) and a strong propensity to self-attract into specialized clusters (Fig. 2c, f), expected from cooperative carotenoid uptake among these proteins (Supplementary Text S1). The transport layer consists of two functional subgroups. The perilipins (PLINs) are maximally disordered, the most efficient drivers of phase-separation, extreme in their avoidance of self-clustering and binding plasticity (Fig. 2), reflecting their roles in sequestration, storage, and distribution of carotenoids (Table S1; Supplementary Text S1). The within-cell transport proteins (FABPs, CRABPs, RBPs) are ordered, weakly phase-separating point- to-point specialist carriers (Fig. 2a-c), although FABP1, FABP4 and FABP5 show extreme binding self-avoidance comparable to PLINs (47-54 × 10⁻³ vs 34-45×10⁻³, correspondingly); CRABP1, CRABP2, FABP2, and FABP6 show moderate self-avoidance (12-23; Supplementary Table S2). This extreme self-avoidance is not an artefact of short length of these proteins (Supplementary Fig. S6) or their IDR span (Methods; Supplementary Text S2; Supplementary Fig. S7b).

**Fig. 2.**
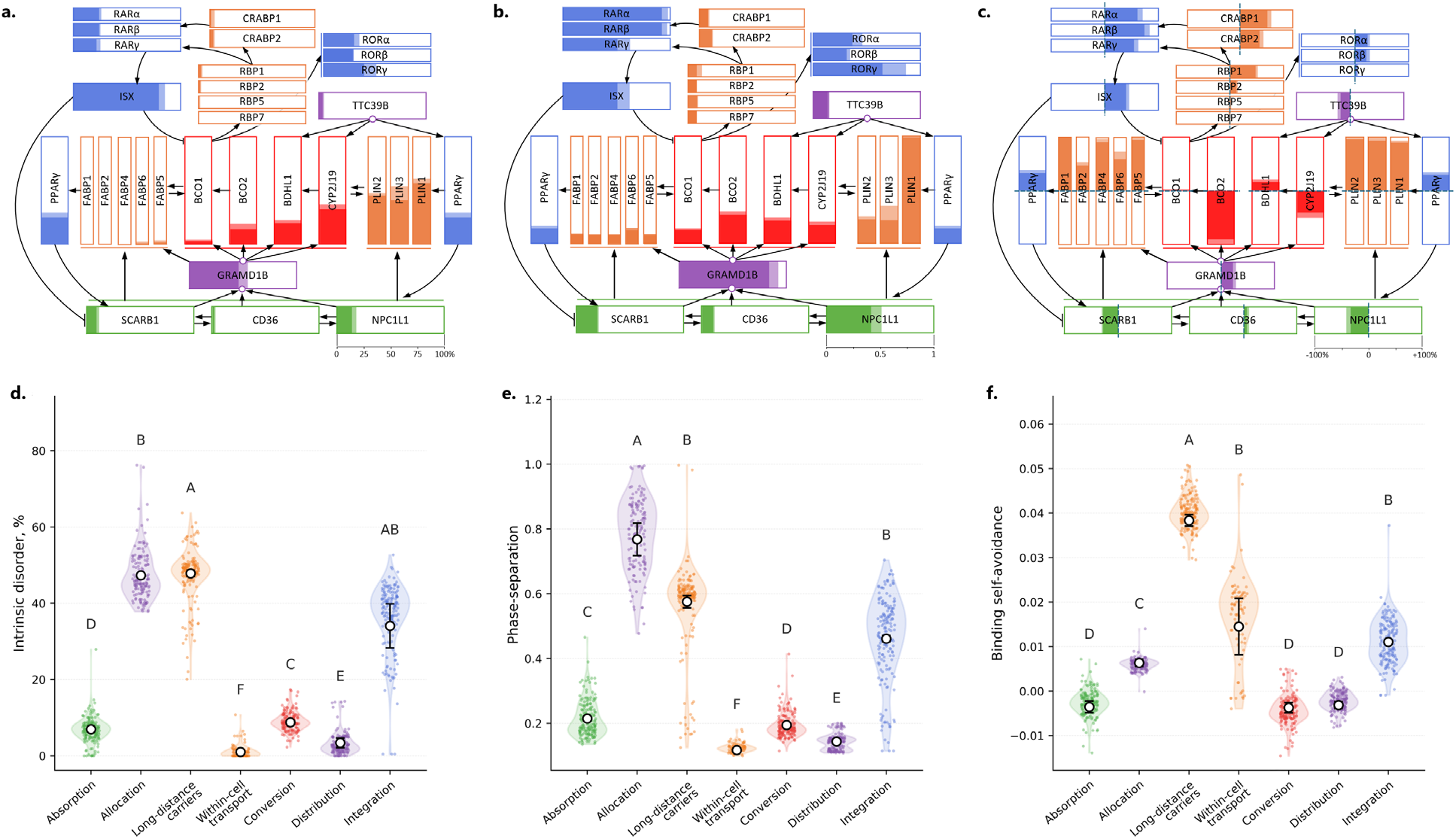
Biophysical profiles reflect functional specialization of proteins in avian carotenogenesis. Colors correspond to functional groups in Fig. 1. The scale for metrics (a-c) is in the lower right corner (on the NPC1L1 bar). Shown are phylogenetic generalized least squares (PGLS) under a Brownian-motion correlation structure with Pagel’s λ estimated from the data (Methods). **(a)** Intrinsic disorder score (% of disordered residues) shown as means (solid color) and standard errors (s.e., semi-transparent color) for each protein in proportion to each rectangle’s area. **(b)** Phase-separation propensity score (ratio, mean + s.e.). **(c)** Homotypic association propensity (standardized by protein length (ε_ST_) and shown as %; negative values indicate binding self-attraction and positive values – binding self-avoidance). **(d-f)** Comparison of biophysical profiles of functional groups. Shown are PGLS estimated means for each functional group ± s.e. computed from the fitted λ-optimized covariance matrix. Transport complex is further partitioned into long-distance and within-cell carriers. Groups sharing a letter are not significantly different at q < 0.05 (the Benjamini-Hochberg correction for multiple comparisons). (d) Intrinsic disorder, (e) Phase-separation propensity, (f) Binding self-avoidance propensity. Based on Supplementary Data S4.

Self-avoidance recurs in other network proteins with connective or regulatory roles. For example, the allocation protein GRAMD1B is highly disordered, strongly phase-separating, and self-repulsive, enabling it to distribute diverse carotenoids from the absorption to conversion modules, whereas the post-conversion distribution protein TTC39B is ordered and weakly phase-separating – reflecting its role as a structured scaffold bridging the integration and conversion modules (Figs. 1b, 2; Supplementary Table S1; Supplementary Text S1). The integration layer shows the connector signature: the transcriptional integrators (RARs, RORs, PPARγ, and especially transcription factor ISX) are highly disordered, phase-separating, and primed for self-avoidant associations (Fig. 2), consistent with their roles in the carotenoid-sensing feedback pathways and transcriptional regulation (Fig. 1b, Supplementary Table S1; Supplementary Text S1). Thus, across the network, the sign of homotypic binding separates two roles – self-attraction of the specialized processors and self-avoidance of the connectors. Strongly self-avoiding allocation and transport layers form a generalized, connective waist in the predicted biophysical bow-tie (Fig. 1c).

### The waist interrupts propagation of specificity enabling network diversification

Tracing homotypic binding along the network pathways reveals that specificity is propagated *within* but is reset *among* the functional complexes by carrier-mediated switches at the waist of the proposed bow-tie (Figs. 1d, 3a). The binding regime changes from self-attracting absorption (phylogenetic mean ε_ST_ = -3.5 × 10^-3^) to self-avoiding allocation (6.4) and transport (33.1), reverting to self-attracting conversion (-3.8) and distribution (-3.1) complexes, and again to self-avoiding integration (11.0), such that every transition in or out of the connector layers is a reset, and every transition between specialized layers is a propagation (Fig. 3a). For example, the self-attracting binding regime propagates unchanged among absorption, conversion and distribution complexes (|Δε_ST_| = 0.2-0.6 × 10^-3^; Supplementary Fig. S2). In contrast, the transport layer is the resetting hub in this architecture: it has the largest cumulative reset than all other layers: 35.9-36.7 against each specialized layer and 21.9-27.3 against the other self-avoiding layers, and it carries 2.2-2.6 times more binding regime transmissions (cumulative ε_ST_ changes) than any other layer, resetting 69% of them (Fig. 3a, Supplementary Data S6). Allocation, by contrast, is the only layer in which propagation exceeds resetting, consistent with its role in transferring carotenoids into the diverse carrier pool. Thus, the overall network is not an array of specialized paths, but a decoupled bow-tie in which self-attracting complexes are linked through a self-avoiding nonspecific waist that consistently interrupts propagation of binding regime between them (Figs.1c, d; 3a). This reset architecture does not depend on whether both long-distance PLINs and within-cell carriers are included in the transport layer (Figs. 2 vs 3). Represented by PLINs alone (the proteins for which the disorder-based model is most appropriate, Supplementary Text S1), the transport layer resets against absorption, conversion and distribution more strongly (Δε_ST_ = 42.0 × 10^-3^) than with all carriers included; represented by only within-cell carriers, the waist similarly strongly resets these layers (18.4 × 10^-3^; *P* < 0.008) (Supplementary Text S2). Nor does it depend on the sequence order: shuffling each sequence while preserving its amino-acid composition reproduces every layer contrast (Supplementary Fig. S8).

**Fig. 3.**
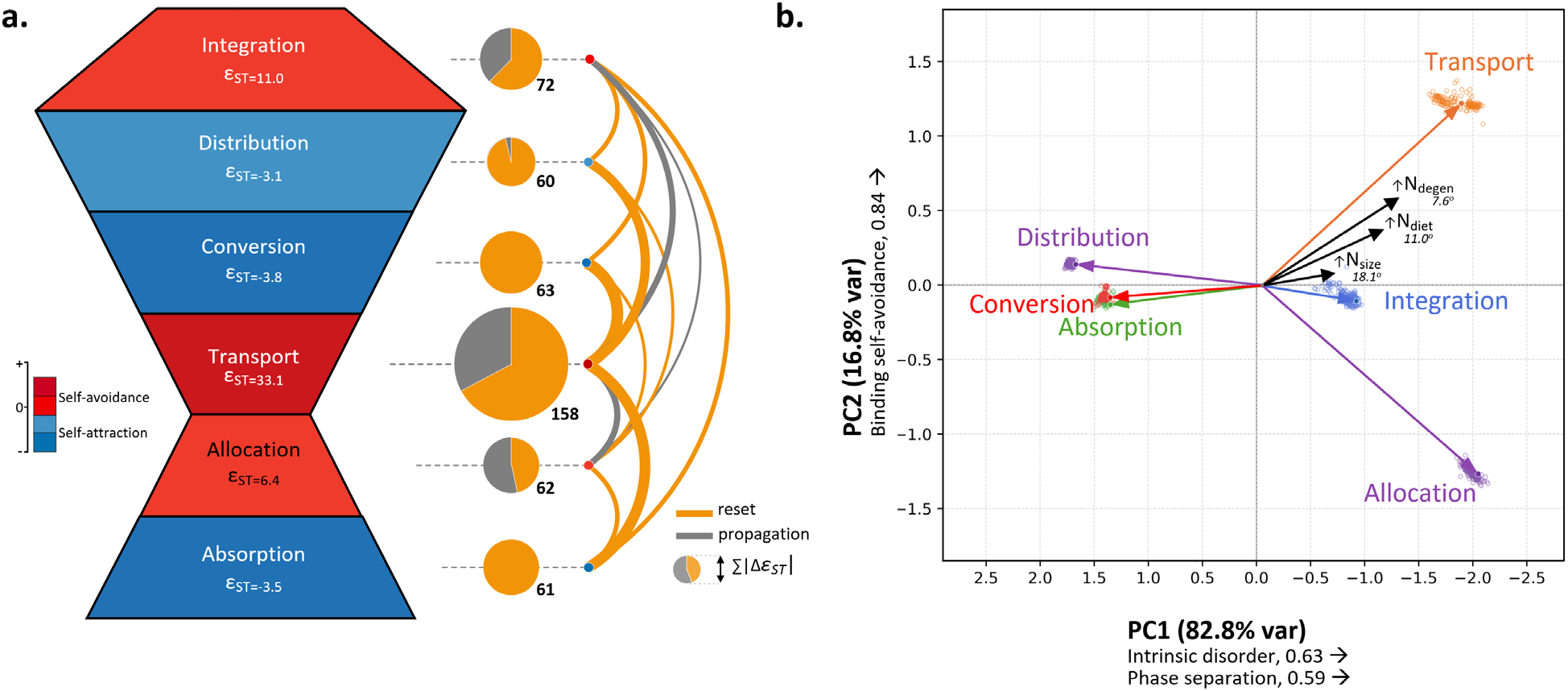
Transport layer consistently interrupts propagation of specialization between functional complexes; its biophysical properties drive network connectivity. (a) Reset architecture of homotypic binding (ε_ST_) across the bow-tie organization of avian carotenogenesis. Layers are stacked in pathway order (Fig. 1b) and colored by the sign of their phylogenetic mean ε_ST_: red, ε_ST_ > 0, binding self-avoidance; blue, ε_ST_ < 0, binding self-attraction). Values within the bow-tie are shown as ×10^-3^. The transport layer combines all storage and transport proteins. Arcs connect layers whose phylogenetic paired contrast in ε_ST_ is significant at *P* < 0.05 (12 of 15 pairs); line width is proportional to |Δε_ST_|; gold arcs mark *resets* (the two layers have opposite binding regimes) and grey arcs mark *propagation* (retention of the same binding regime). Circle area is proportional to Σ|Δε_ST_| over the layer’s links with the share of resets (gold) versus propagation (grey), based on Supplementary Fig. S2. (b) Functional layers on the principal component (PC) plane of the three disorder properties calculated on phylogenetically transformed values. Points are species values, solid dots are centroids, and lines connect centroids to the origin. Black vectors show the direction of increase of network size (N_size_, PC1 of the number of nodes and edges), number of degenerate nodes (N_degen_) and number of dietary carotenoids (N_diet_) in species networks, projected on the same plane; lengths are proportional to their correlation with the transport vector, angles are relative to the transport vector. All three network measures increase toward greater disorder, phase separation and binding self-avoidance of the transport protein group. Based on Supplementary Data S5.

Across species, larger, more diverse (greater number of dietary carotenoids) and more degenerate networks are most closely associated with greater disorder, phase-separation and self-avoidance of their carrier proteins but not with biophysical properties of their enzymatic or absorption complexes (Figs. 3b, 4b). Further, partitioning the variance in each network measure among the three biophysical properties within each layer shows that the two self-avoiding layers of allocation and transport, comprising the waist, account for 22-29% of the among-species variance in network size and degeneracy and 16% of the variance in the number of dietary carotenoids for transport, whereas absorption, conversion and distribution complexes do not explain any variance in network parameters (Fig. 4a, b). Most of the effect of transport proteins, and the effect of allocation layer on network size, is attributable to ε_ST_ (Fig. 4a), identifying binding self-avoidance as the primary driver of network growth (Supplementary Figs. S4 and S5). These results support the model of network branching after resets. For example, transport-conversion reset affects network size overwhelmingly through the branching of degenerate paths downstream of the reset rather than the addition of dedicated diet-specific paths (Fig. 4c).

**Fig. 4.**
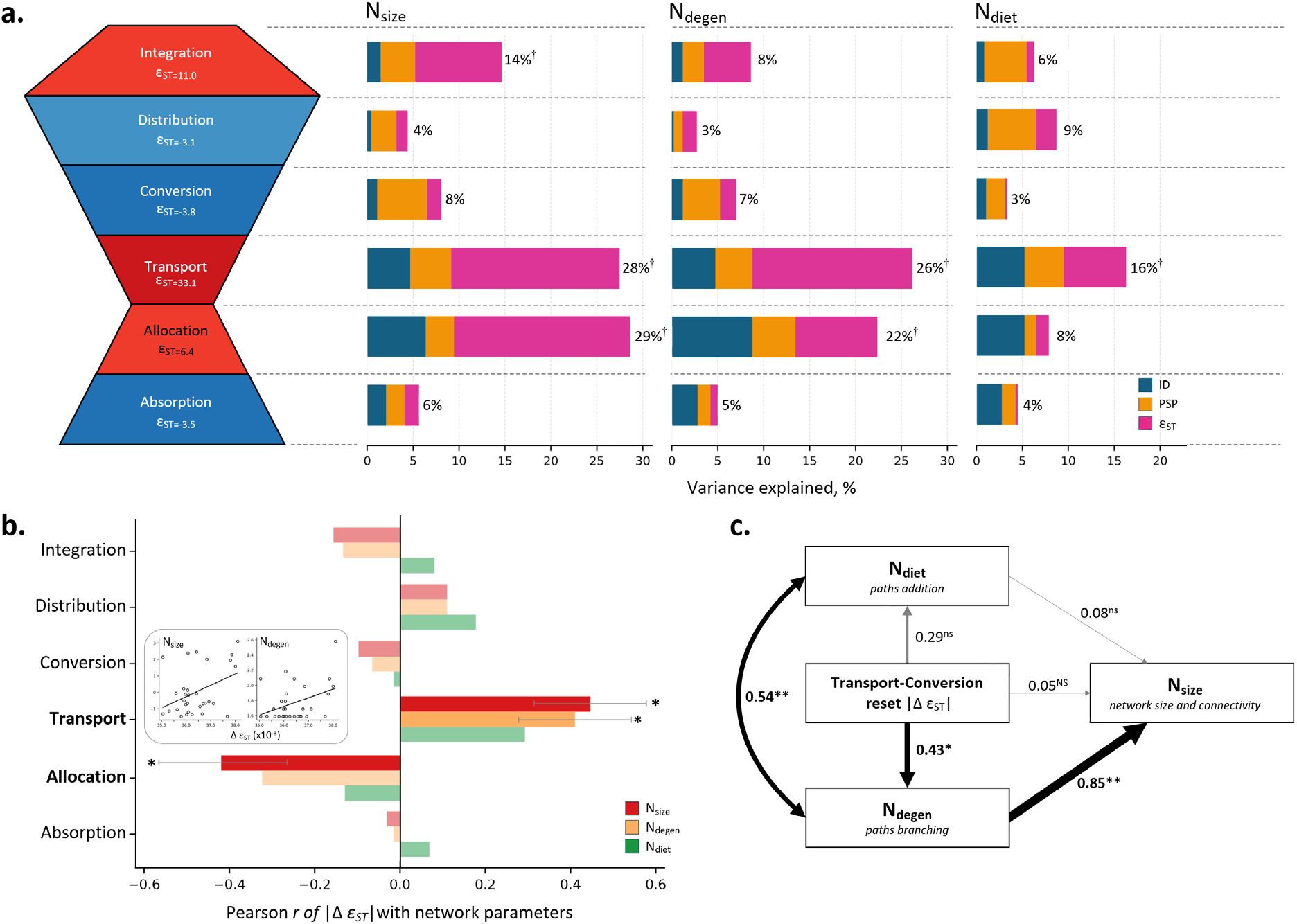
Binding self-avoidance of the waist proteins drives network growth and connectivity by interrupting propagation of specialization and inducing post-reset pathway branching. (a) Among-species variance (%) in network size (N_size_), number of degenerate nodes (N_degen_) and number of dietary carotenoids (N_diet_) explained by the intrinsic disorder (ID, blue), phase-separation propensity (PSP, orange) and homotypic binding (ε_ST_, pink) of each functional layer, from multiple regressions on phylogenetically transformed values (Supplementary Fig. S3, Supplementary Data S6); stacked bars are the incremental decomposition of *R^2^* (Methods) aligned to the bow-tie layers at left (layer colors and ε_ST_ as in Fig. 3a; †*p* < 0.1). Supplementary Figs. S4 and S5 show contributions while accounting for collinearity of the three disorder measures. (b) Species whose networks have more self-avoiding transport proteins and more self-attracting allocation proteins have larger and more interconnected networks. Insets show partial regression plots of N_size_ and N_degen_ on the magnitude of the waist reset, |Δε_ST_|. Shown are summary Pearson *r*’s per layer on phylogenetically transformed values; whiskers are 95% confidence intervals for correlations significant at \**p* < 0.05. Non-significant correlations are shown as half-transparent bars. Supplementary Fig. S5b shows all transitions within each layer. (c) Path model (b_st_ coefficients on phylogenetically transformed values) for a subset of *n* = 31 species linking the transport-conversion reset magnitude |Δε_ST_| to network size (PC1 of number of edges and number of nodes) through the two routes of network growth predicted in Fig.1e: addition of dietary input paths (N_diet_) or branching of degenerate paths (N_degen_). Line width is proportional to b_st_ (\**p* < 0.05, \*\**p* < 0.01, *ns*, not significant). The effect of the transport-conversion waist reset on network growth is carried almost entirely by branching (indirect effect through N_degen_ ≈ 85% of the total), as expected if resets decouple upstream origin from downstream recombination (Fig. 1e).

## Discussion

Taken together, these results show how a biophysical bow-tie architecture converts external diversity into internal precision in avian carotenogenesis. First, specialized complexes of dietary absorption and enzymatic conversion are ordered, weakly phase-separating, and self-attracting, whereas the allocation and carrier proteins that connect them are disordered, phase-separating, and strongly self-avoiding (Fig. 2). Second, this biophysical contrast consistently resets, rather than propagates, binding specificity across the network (Fig. 3a), with transport proteins being the principal hub at which input identity is decoupled from downstream utilization. Third, across species, network diversification is closely associated with the magnitude of this waist reset but not with the properties of the absorption or conversion complexes (Figs. 3b, 4); this association is specifically produced by the accumulation of degenerate branching, rather than the addition of dedicated diet-carotenoid-specific paths (Fig. 4c). These findings have three evolutionary implications.

The first is that the connections among conserved modules rather than the modules themselves harbor the greatest potential for innovation and diversification. Much work on metabolic evolvability emphasizes catalytic promiscuity, gene duplication, and changes in substrate specificity (41, 42). In avian carotenoid networks, however, evolutionary diversification proceeds by changing the routing among conserved (43, 44) modules rather than by changing their catalytic identity: species with larger and more diverse networks differ most strongly in the biophysical properties of the connector layers that reset the specificity of inputs, enabling them to be routed between multiple conversion modules (Fig. 5a) (see also 14 for experimental illustration that reorganizing interactions among conserved enzymes redirects metabolic flux without changing catalytic specificities). This suggests a mechanistic basis for the often-documented combinatorial evolution, flux-structure concordance, weak phylogenetic dependence, and recurrent reuse and introgression of conserved enzymatic machinery in avian carotenoid evolution (31, 37, 40, 45–48).

**Fig. 5.**
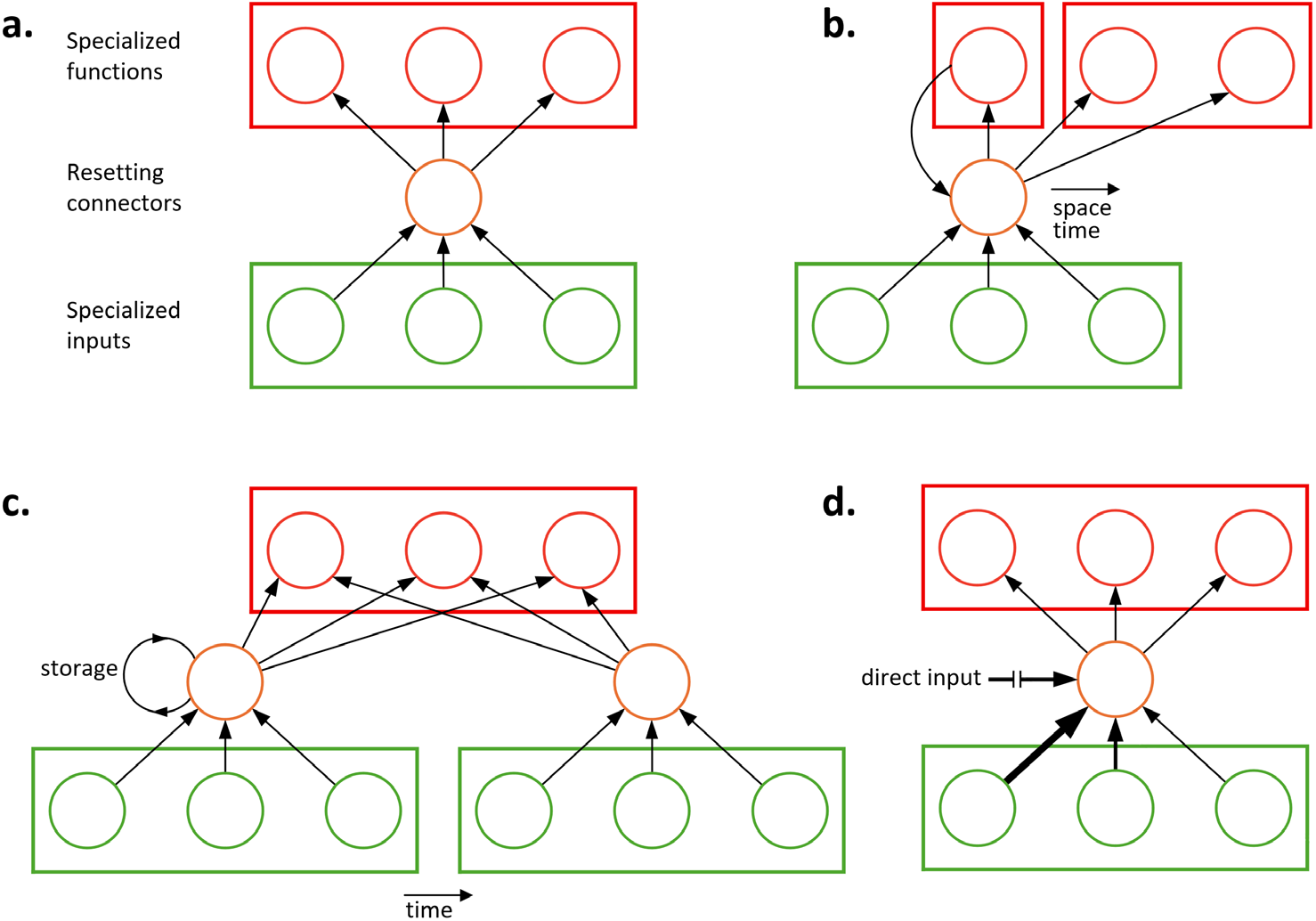
The biophysical bow-tie organization. **(a)** where specialized layers of acquisition of external carotenoids (green) and of their integration into specialized internal functions (red) are linked by connectors non-specific to either of these functions (orange), meets two essential requirements of avian carotenogenesis: openness to novel dietary carotenoids and the need to accumulate metabolic degeneracy to sustain their internal integration. It may also account for central phenomena in avian carotenoid evolution: **(b)** The ubiquitous recombination of enzymatic modules where non-specific carriers can exchange, recombine, and sequentially link substrates among conversion modules separated in space and time, resulting in internal **elaboration and diversification** of carotenoid pathways, **(c)** the ability to **amplify and direct** carotenoid flux in a time-and tissue-specific manner facilitated by the carrier’s propensity to store, aggregate, and redistribute dietary carotenoids, and **(d)** notable **openness to lateral transfers** of substrates that predisposes carotenoid networks to symbiotic associations and introgressed transfers of enzymatic complexes, capitalizing on the carriers’ ability to transfer products not only by protein binding studied here (arrow thickness indicate different substrate affinities), but also by indiscriminate lipid-droplet enclosure (18) (Supplementary Text S1).

The second implication is that external dependence does not necessitate external control. Birds are obligatorily dependent on dietary carotenoids, yet it is the nonspecialized carrier layer that determines how these external inputs are stored, combined and redirected to downstream specialized modules. Theory distinguishes between source controls imposed at a network’s external entry points and internal control generated by branching expansion: when upstream states branch into a larger set of downstream states, additional internal controls are required for network functionality (49–51). We previously showed that gains of redundant dietary inputs in avian carotenoid networks produce shifts from source-controlled to internally-controlled metabolic networks and that evolutionary cycles in these control shifts are reflected in the tempo and mode of avian carotenoid diversification (32). For example, acquisition of internal controls occurred in less than 9% of phylogenetic transitions of control profiles of biochemical networks across 250 avian species yet accounted for nearly 70% of variation in evolutionary rates of carotenoid pathways because it enabled sustainable exploration of internal metabolic branching. The current study provides a molecular basis for these patterns: self-avoiding carriers reset input-specific interactions, allow substrates to be recombined between specialized conversion modules, leading to accumulation of degenerate branching (Fig. 4c), which, in turn, is strongly linked to gaining greater internal control (N_degen_ vs number of internally controlled nodes (N_i_), *b*_ST_ = 0.88, *n* = 243 species, *P* < 0.001) (32) rather than favoring elongation of dedicated input-output pathways that would have retained their source controls (40). The biophysical bow-tie waist is thus the link between redundancy of dietary carotenoids and the internal control of their utilization, whereas non-specificity of the waist proteins is the mechanism that buffers the contingency of external inputs from the precision of internal integration.

Finally, the distinction between external dependency and network control accounts for several key features of avian carotenogenesis (Fig. 5b-d). Because carriers are not carotenoid-specific and yet closely integrated with organismal condition and physiological state (52, 53), conserved enzymatic modules are commonly redeployed across tissues, developmental stages, sexes, and seasons without modifying the modules themselves (Fig. 5b)(54–57). Storage, aggregation, and redistribution of substrates by the carrier layer can also amplify or redirect flux to particular tissues, life-history stages, or physiological states (e.g., during oogenesis, molt, or during starvation) (34, 58–61) (Fig. 5c), again without modification of the conversion machinery. This organization may also account for notable priming effects in which exposure to distinct dietary carotenoids across successive molts or reproductive episodes increases the efficiency of the same downstream pathways, resulting in “emergent buffering” of internal functions from volatility of dietary inputs (Fig. 5c)(62, 63). The same architecture also enables laterally acquired “foreign” inputs to enter established downstream pathways (Fig. 5d). Carotenoids supplied through direct transfer (27, 64–67), microbial metabolism or symbiosis (68–70), or by introgressed enzymatic modules (46–48) can be accommodated precisely because the waist weakens the dependence of downstream routing on upstream origin. Thus, the three most defining properties of avian carotenogenesis – physiological sensitivity (“condition-dependence”), openness to novel or laterally acquired inputs, and the evolution of greater internal metabolic control – can all be understood as consequences of the same architecture: decoupling of external origin from internal integration enabled by nonspecific intermediate connectors in the bow-tie organization.

More generally, our results suggest that the precision of a biochemical outcome need not reflect the specificity of each preceding molecular interaction. In the proposed bow-tie architecture, specialized modules retain their catalytic constraints, whereas a carrier-rich waist limits the propagation of input-specific routing. Downstream outcomes may therefore depend on the organization of accessible functional modules and their collective interactions, rather than on dedicated recognition at each intermediate step. This principle parallels the emergence of specificity in transcription factor localization from ensembles of individually nonspecific interactions (9, 71). Importantly, the same type of interactions can support distinct physical outcomes: transient scaffolding may favor selective recruitment, whereas phase transition can disrupt that recruitment and favor sequestration. In carotenoid networks, the balance between exchange and compartmentalization could permit buffering of fluctuating inputs without fixing their downstream fate (Fig. 5). Direct measurements of carrier–cargo interactions, substrate flux, and compartmentalization will be required to test this proposed mechanism.

These findings also suggest a dynamic sequence by which strict dependency on unpredictable external resources is reconciled with sustained internal precision and evolvability – a ubiquitous combination in organic evolution. Specialized modules preserve precision; self-avoiding connectors compartmentalize that specificity, allowing modules to be recombined while maintaining network cohesion; recombination generates degeneracy of connectors that shifts network control from external sources towards internally driven dynamics; interdependence of external input redundancy and internal control resets the cycle. The ability to interrupt the propagation of specialization may therefore be as important for evolutionary diversification as the ability to generate specialization itself. Illustrated here by a particularly tractable example of avian carotenoid evolution, this process might reflect a fundamental principle by which gains and losses of external dependencies drive dynamic evolution.

## Materials and Methods

### Protein network of avian carotenogenesis

Fig. 1b and Table S1 summarize functional groups of 31 core proteins of avian carotenogenesis and their interactions. Supplementary Text S1 provides the empirical basis for this summary.

### Protein sequences and protein disorder analyses

Amino acid sequences for 31 avian carotenogenesis proteins were obtained for 172 species (Supplementary Data S1, S1a, S2) following quality-filtering protocols and scripts described previously (19). Special care was taken to retain only the full-length (>90%) sequences, which is particularly important for enzymes and other membrane-embedded proteins in this dataset (e.g., NPC1L1, SCARB1, CD36) because, for these proteins, it is common in the literature to truncate the protein length at the membrane anchor. After quality filtering, for each orthologous protein sequence, we quantified three sequence-derived biophysical properties that predict (i) intrinsic disorder, (ii) droplet-forming phase-separation propensity, and (iii) the net strength and sign of chemically specific homotypic interactions that promote self-association or avoidance. We averaged these values across all isoforms for each protein within a species (Supplementary Data S1).

Intrinsic disorder parameters were calculated with MetaPredict (https://metapredict.net/), a machine-learning predictor trained to reproduce consensus disorder scores derived from several established disorder predictors (72, 73). We computed the protein-level fraction of disordered residues as the percentage of residues with MetaPredict disorder score above a disorder threshold of 0.5. To summarize intrinsically disordered region (IDR) architecture, we identified all contiguous segments of residues with MetaPredict disorder score > 0.5 and recorded the longest IDR per protein (Supplementary Data S1). Predicted phase-separation propensity was quantified with FuzDrop (https://fuzdrop.bio.unipd.it/predictor), a sequence-based method that predicts the probability of spontaneous liquid-liquid phase separation (LLPS) and identifies droplet-promoting regions (74). The approach is based on a model in which the droplet state is stabilized by disordered interactions; the method combines predictions of disordered binding modes (FuzPred) with structural disorder estimates (Espritz) and integrates these features into protein-level droplet propensity predictions. Proteins with pLLPS ≥ 0.60 were considered phase-separation drivers (74, 75).

To measure protein interactions, we used FINCHES (https://github.com/idptools/finches), a sequence-based framework that capitalizes on physicochemical potentials developed for molecular simulations to predict intermolecular interactions driven by IDRs (39, 76, 77). The metric estimates residue–residue interaction preferences and summarizes net interaction strength as a homotypic interaction parameter (ε), which reflects the balance of attractive (negative ε) vs. repulsive (positive ε) interactions in the mean-field limit. We computed protein-level ε and then standardized ε by protein length to obtain a per-residue interaction density ε_ST_ that is comparable across the variable-length proteins under this study (Supplementary Data S1); ε_ST_ is the mean pairwise interaction parameter of the sequence that does not depend on its length (Supplementary Text S2). Because FINCHES models chemically specific, disorder-mediated interactions and does not capture structured, sequence-specific interfaces, ε_ST_ values for the largely folded specialized layers are used here as qualitative proxies for the disorder-mediated component of their surface chemistry, with specific focus on *relative* differences between proteins in the network (Fig. 2; Supplementary Text S2). The same applies to the compact β-barrel within-cell carriers (FABPs, CRABPs and RBPs). To establish the suitability of this approach, we explicitly tested the robustness of ε_ST_ to variation in (i) protein length, net charge per residue and aromatic content across proteins, and recomputing it for tandemly duplicated sequences and for 132-residue windows of longer proteins; (ii) recomputing ε over IDRs only (MetaPredict score ≥ 0.5), folded residues only, and residues with relative solvent accessibility ≥ 25% (DSSP), using experimental structures of mammalian orthologues of the 11 within-cell carriers and CD36 and of paralogues of SCARB1 (CD36) and of BCO1 and BCO2 (RPE65); (iii) comparing each observed ε_ST_ with composition-preserving reshuffles of the same sequence (Supplementary Text S2, Supplementary Tables S2-S4, Supplementary Data S7). An important next step is direct empirical assessment of the extent to which sequence-based predictions reflect biophysical properties.

### Phylogeny and species datasets

A time-calibrated maximum clade credibility global avian phylogeny of 11,145 species based on the 2024 eBird/Clements Taxonomy (The Aves tree v1.5) (78) was pruned to 172 species included in this study (Supplementary Fig. S1, Supplementary Codes S1, S2) using the R package clootl 0.1.2 (79). Carotenoid network data for the species under this study (Supplementary Data S1, S1a) are from a comprehensive dataset of avian carotenoid networks (80).

### Statistical analyses

We compared intrinsic-disorder metrics among functional groups using phylogenetic generalized least squares (PGLS) under a Brownian-motion correlation structure with Pagel’s λ estimated from the data, based on time-calibrated phylogeny (Supplementary Fig. S1). For each species, we computed the protein-level metrics across all proteins assigned to one of the six functional groups (Absorption, Allocation, Transport, Conversion, Distribution, and Integration, Fig. 1). Only species with complete data across all functional groups for a given metric were retained so that comparisons involving all groups used an identical species set. In Fig. 2a-c we report analyses for all individual proteins; two proteins that had low species coverage after the quality control – CYP2J19 and PPARγ (Supplementary Data S1) – were excluded from the functional *group* level analyses. To normalize distributions and stabilize variance, we log10-transformed MetaPred% and FuzDrop measures and arcsinh-transformed ε_ST_.

To compare differences among functional groups (Fig. 2), we performed pairwise comparisons using phylogenetically corrected contrasts by fitting PGLS models to within-species group differences and testing whether the phylogeny-adjusted mean difference differed from zero with Benjamini– Hochberg false-discovery-rate correction (with FDR q < 0.05) to adjust for multiple comparisons (Supplementary Data S4). To visualize functional group separation in multivariate disorder space, we first computed phylogenetically transformed values for each functional group using λ optimized for that group (Supplementary Data S4). All transformed observations were then pooled across functional groups and standardized (z-scored using the global mean and s.d.) and projected onto a common principal component (PC) plane based on three disorder measures (Supplementary Data S5). PCs were obtained by singular value decomposition of the standardized transformed matrix.

Network measures (number of nodes, NodeN; number of edges, EdgesN; number of dietary carotenoids, N_diet_; number of degenerate nodes, N_degen_) were phylogenetically transformed with the same λ-optimized PGLS procedure used for the protein metrics (Supplementary Data S5). Network size (N_size_) was PC1 of a principal component analysis of the standardized, phylogenetically transformed NodeN and EdgesN; PC1 loaded equally on both variables (∼0.71) and explained 98.7% of their variation (eigenvalue = 1.98). For projection of network measures onto the disorder PC plane (Fig. 3b), we plotted one observation per species (the mean of its group-centered PC scores). The direction of increase of each network measure was obtained from its correlations with the PC1 and PC2 coordinates across species; vector angles were relative to the transport group vector, and vector lengths were scaled to that correlation. To quantify associations between disorder and carotenoid network properties within functional groups, we fitted, separately for each group, multiple regressions of each network measure on the group’s three phylogenetically transformed properties (ID, PSP, ε_ST_) and, in a parallel analysis, on the first two principal components of a within-layer PCA of the same three properties (Supplementary Data S6; Supplementary Figs. S3 and S4).

### Phylogenetic analyses of ε_ST_ resets and network evolution

To test whether the binding regime propagates or is reset between functional complexes (Fig. 1d), we calculated, for each species, the mean ε_ST_ of the proteins of each functional group. For each group, we estimated a phylogenetic mean by GLS under the Brownian covariance matrix of the pruned tree (Supplementary Data S2a), with Pagel’s λ estimated by maximum likelihood; these GLS intercepts are the group ε_ST_ values reported in Figs. 3a and 4a, and their sign defines the group’s binding regime. Between-layer resets were tested with phylogenetic paired contrasts: for each of the 15 layer pairs, we calculated the within-species difference in ε_ST_ between groups and fitted a λ-optimized GLS model to these differences, yielding the phylogeny-adjusted mean reset Δε_ST_, its s.e. and a *t*-test of Δε_ST_ = 0 (Supplementary Data S6). We then correlated group ε_ST_ values and reset magnitudes with the phylogenetically transformed network measures (Pearson *r*, 95% CI from Fisher’s *z* transformation). The relative contribution of ID, PSP and ε_ST_ to variance in each network measure within each layer (Fig. 4a, Supplementary Fig. S5) was quantified from the multiple regressions by the variance decomposition, which averages the sequential increment in *R*^2^ of each predictor over all orderings of the predictors (the Lindeman-Merenda-Gold decomposition (81)) and yields non-negative shares that sum to the model *R*^2^; this decomposition was used because the three properties correlate within layers (Supplementary Fig. S5).

The two routes of network growth predicted in Fig. 1e were compared with a path model (Fig. 4c) on standardized, phylogenetically transformed variables for a subset of 31 species with complete network data and transport-conversion reset data. Total, direct and indirect effects were obtained as products of standardized path coefficients with PROC CALIS (SAS 9.4); the sampling distributions of the indirect effects were examined by non-parametric bootstrap (10,000 resamples of species).

## Supplementary Information

**Supplementary Data S1:** Biophysical properties of 31 core carotenogenesis proteins across 172 species and network data

**Supplementary Data S1a:** Summary of S1 by functional groups of Fig. 1b.

**Supplementary Data S2:** Phylogeny for 172 species and phylogeny references

**Supplementary Data S2a:** Brownian covariance matrix for PGLS (Figs. 2–4) showing shared root-to-MRCA branch length for every pair of species (diagonal = root-to-tip depth)

**Supplementary Data S3:** Amino acid sequences for 31 proteins for 172 study species (FASTAS.zip)

**Supplementary Data S4:** Raw and phylogenetically transformed biophysical properties by functional group for Figure 2.

**Supplementary Data S5**: Principal-component analysis of phylogenetically transformed biophysical properties and projection of carotenoid network measures for Figure 3b

**Supplementary Data S6:** Phylogenetic analyses of ε_ST_ resets and their association with network parameters (Figs. 3a, 4; Supplementary Fig. S2–S5)

**Supplementary Data S7:** Results of robustness tests (Supplementary Text S2; Supplementary Tables S2–S4).

**Supplementary Code S1-S6:** Code for protein pipeline, phylogenetic reconstruction, and robustness tests

**References to Supplementary Information**

### Supplementary Text S1

#### Protein network of avian carotenogenesis

Figure 1b and Supplementary Table S1 summarize functional groups of 31 core proteins of avian carotenogenesis (Supplementary Data S1) and their interactions. Below is the empirical basis for this summary.

##### DIETARY ABSORPTION GROUP (SCARB1, CD36, NPC1L1)

Three membrane proteins on the intestinal brush border facilitate uptake of carotenoids from micelles into enterocytes. After uptake, absorbed carotenoids are incorporated into portomicrons and carried to the liver and other tissues via the portal circulation (1, 2).

###### SCARB1

An evolutionarily and functionally conserved cell-surface HDL receptor that mediates the selective uptake of carotenoids (3, 4). SCARB1 is highly expressed on enterocyte microvilli, where it directly binds carotenoid-rich micelles and transports carotenoids into the cell (5–7). Wild-type SCARB1 promotes carotenoid absorption, whereas the loss-of-function mutant leads to carotenoid deficiency in pigmentation, although additional pathways of carotenoid absorption (below) or passive diffusion can partially compensate (8, 9). In addition to the intestine, SCARB1 is expressed in the liver and retina, where it imports carotenoids from circulating lipoproteins (3, 4, 10) and mediates uptake of dietary xanthophylls (11). Colocalization at the brush border facilitates cooperation between SCARB1 and CD36 (5). SCARB1 also forms close functional associations with downstream metabolic enzymes (Fig. 1b) and transporters inside the enterocyte. For example, in intestinal cells, absorbed β-carotene from SCARB1 is immediately cleaved by BCO1 into retinaldehyde (12, 13). The resulting retinaldehyde is then bound by cellular retinol-binding protein (RBP2, see below) in the same cells (14), confirming a close functional coupling of Absorption and Conversion group (Fig. 1b). SCARB1 is also subject to feedback regulation by vitamin A levels; once dietary provitamin A carotenoids have been converted to retinoic acid (RA), RA activates retinoic acid receptor (RAR) in the intestinal cells, which induces ISX (see Integration, below) to repress the expression of both SCARB1 and BCO1 (15, 16) (Fig. 1b).

###### CD36

A scavenger receptor on enterocyte brush-border membranes that binds a broad range of hydrophobic molecules, including carotenoids (5, 17). In adipose or immune cells with low concentrations of circulating carotenoids, CD36 variation is associated with carotenoid absorption, contributing to integument pigmentation (18). Similarly to SCARB1, it forms a regulatory feedback loop with vitamin A accumulation and conversion: a SNP upstream of CD36 alters RAR binding sites; the allele with higher promoter activity is associated with more pigmented skin (18). Cooperation of CD36 and SCARB1 is further reinforced by coordinated regulation by RA-ISX feedback (16). Outside the gut, CD36 is broadly expressed (e.g., in liver and retina) and may mediate tissue-specific carotenoid uptake (Supplementary Table S1) (5, 17).

###### NPC1L1

A transmembrane transporter located on the apical surface of enterocytes that facilitates carotenoid uptake in the intestine using a vesicular endocytic mechanism to internalize sterols (with carotenoids) from the gut lumen (7, 19). Experimental inhibition of NPC1L1 reduces intestinal uptake of β-carotene and α-carotene more than that of lutein and zeaxanthin (7), suggesting selective absorption of carotenoids across the three absorption proteins studied here. In chickens, NPC1L1 is functionally integrated with SCARB1 and CD36 in the gut and the liver (20), showing coordinated response to selection for greater carotenoid absorption and pronounced compensatory interactions (above), despite differences in the absorption mechanisms. Similar to other genes in this functional group, NPC1L1 is regulated by the ISX-mediated feedback. Additionally, NPC1L1 on hepatocyte membranes of the liver may direct circulating carotenoids to storage (Fig. 1, see Transport module, below).

##### CHEMICAL CONVERSION GROUP (BCO1, BCO2, CYP2J19, BDH1L)

Two major groups of enzymes carry out chemical modifications in birds: carotenoid cleavage dioxygenases (which cleave carotenoids to smaller apocarotenoids or vitamin A compounds) and carotenoid ketolases/dehydrogenases (which add oxygen functional groups, producing modified pigments) (21–24). The key avian enzymes in these categories are BCO1 and BCO2 (carotenoid oxygenases), and CYP2J19 together with BDH1L (ketolation pathway enzymes). Their expression is often tissue-specific: BCO1 dominates vitamin A-producing tissues, while CYP2J19/BDH1L predominate in red-pigment-producing tissues.

###### BCO1

A cytosolic enzyme that symmetrically cleaves β-carotene (and other provitamin A carotenoids) at the central 15,15′ double bond to yield two molecules of all-trans retinal (retinaldehyde) (12, 24, 25). In birds, BCO1 is highly expressed in the intestinal mucosa and other tissues that generate retinaldehyde for vision (e.g., the retina) or storage (e.g., the liver) (13, 25). The immediate product of BCO1, retinaldehyde, can be reduced to retinol or oxidized to RA to regulate gene expression (24, 26). BCO1 can operate immediately downstream of the dietary uptake step in provitamin A metabolism. For example, in an intestinal cell, dietary β-carotene is taken up via SCARB1, then BCO1 cleaves it to retinaldehyde, which is immediately bound by cellular retinol-binding protein (RBP2; Transport group, below; Fig. 1b) (14, 27). This close coupling and capture by RBP2 prevents retinaldehyde from diffusing or causing oxidative damage. BCO1 is functionally coexpressed with both SCARB1 and RBPs (8, 14). The two enzymatic modules in the chemical conversion group (Fig. 1), BCO1 and CYP2J19, are physically and physiologically separated (28–30). Tissues specialized in pigmentation tend to have low BCO1 expression, presumably to avoid diverting carotenoids into vitamin A pathway (28). Intestinal BCO1 is also repressed by RA-induced ISX, similarly to SCARB1, to limit vitamin A excess, with experimental confirmation of this regulation in mammals (15, 16). Birds rely on BCO1 for obtaining vitamin A from diet, so variations in BCO1 expression can influence how much carotenoid is available for coloration versus conversion; the allocation is likely regulated by downstream genes in the Integration group (see below).

###### BCO2

A mitochondrial enzyme that asymmetrically cleaves carotenoids at alternative double bonds, typically cleaving one β-ionone ring from the molecule. BCO2 has a substrate preference for xanthophylls (24, 31). Functionally, BCO2 serves as a carotenoid disposal or tuning pathway: by cleaving carotenoids, it prevents excessive accumulation and can generate signaling molecules from carotenoids. For example, in mice, BCO2 deletion leads to high xanthophyll accumulation in tissues and associated mitochondrial oxidative stress (32). Tissues that strongly deposit carotenoids often downregulate BCO2, facilitating accumulation (32, 33). Conversely, in fish, tissues that remove carotenoids upregulate BCO2 facilitating their breakdown (34, 35). BCO2 and BCO1 can act sequentially (Fig. 1b): BCO2’s initial cleavage yields an apocarotenal that can then be further cleaved by BCO1 to retinal (31, 36). This two-step route allows metabolism of carotenoids that BCO1 alone cannot efficiently cleave. However, because BCO2 resides in mitochondria and BCO1 in the cytosol, any shared substrates must either diffuse or be transported (see Transport group, below) (23, 32). BCO2’s pathway is functionally isolated from the ketocarotenoid pathway because coexpression of BCO2 in a cell that is converting red ketocarotenoids via CYP2J19 would destroy the substrate (32). BCO2 is coexpressed with GRAMD1B (Fig. 1b; Allocation group, below). GRAMD1B can allocate and deliver xanthophylls to the mitochondrial membrane for BCO2 (37), such that carotenoids allocated for degradation are funneled into mitochondria. BCO2 expression is also influenced by the Absorption functional group (see above).

###### CYP2J19

A cytochrome P450 enzyme that, working with the dehydrogenase BDH1L, converts yellow dietary carotenoids into red C4-ketocarotenoids (29, 30). CYP2J19 is expressed in the dermis of growing pigmented feathers, in the liver, and in the red cone photoreceptors of the retina. First, CYP2J19 hydroxylates C4 (and C4′) of the β-ionone rings of dietary carotenoids, producing a 4-hydroxycarotenoid intermediate (22, 28). Then, BDH1L oxidizes this C4-hydroxyl to a ketone, giving the red C4-ketocarotenoid. Any 3-hydroxyls already on the ring (as in zeaxanthin) are kept, so zeaxanthin is converted to astaxanthin. Thus, CYP2J19 and BDH1L function as a sequential pair, producing the red pigments from dietary precursors (28, 38) (Fig. 1b), and are typically colocalized in the same cells. CYP2J19 may not directly interact with retinoid-binding proteins (RBPs, below); it is located in the endoplasmic reticulum (ER) and works on intact carotenoids, so its product (ketocarotenoid) does not significantly contribute to RA levels (39). TTC39B (Distribution group, below) is an accessory factor coexpressed with CYP2J19. In birds and fish, TTC39B expression levels modulate how efficiently ketocarotenoids are produced (28, 40, 41), possibly by affecting enzyme stability (see below).

###### BDH1L

A short-chain dehydrogenase/reductase which is the partner enzyme to CYP2J19. In the red pathway it oxidises the C4-hydroxyl introduced by CYP2J19 to a C4-keto group (28, 42). Acting alone, BDH1L instead converts the 3-hydroxy-β-rings of zeaxanthin or lutein into yellow ε,ε-ketocarotenoids; for instance, in canaries, BDH1L alone converts zeaxanthin to canary xanthophyll B (22, 28). BDH1L acts sequentially after CYP2J19; it likely accepts the hydroxylated carotenoid released by CYP2J19 in the vicinity of the ER membrane. Association between BDH1L and CYP2J19 enzyme complex might be maintained by intracellular transport as both enzymes are membrane-associated (39, 42).

##### Allocation group

###### GRAMD1B

Segregates carotenoids into the appropriate compartment pre-modification (Fig. 1b). GRAMD1B is a member of the Aster protein family that mediates non-vesicular lipid transport between cellular membranes (43, 44). It contains a GRAM domain that allows it to tether to the plasma membrane and a StARkin domain that binds cholesterol and similar lipids (44). In mammals, GRAMD1B ferries cholesterol from the plasma membrane to the ER for homeostatic regulation (44). It can also bind and transport xanthophylls inside cells to specific locations (37). GRAMD1B is thought to function at the interface of the membrane and cytosol to partition carotenoids: e.g., directing carotenoids to the mitochondria for BCO2, or to the ER for incorporation into lipoproteins or droplets (37, 45). GRAMD1B is broadly expressed because of its involvement in cholesterol trafficking; thus, it is likely available in most cells that handle carotenoids. GRAMD1B is functionally coexpressed with BCO2 (above) and by affecting the availability of carotenoids (or their metabolites) in the cell, it influences linked pathways through regulation by proteins in the Integration group (Fig. 1b), or by inducing allocation trade-offs between tissues in organelle-specific delivery (24).

##### Distribution group

###### TTC39B

Enhances a specific metabolic outcome by post-conversion allocation (Fig. 1b, Supplementary Table S1). Closely coexpressed with carotenoid ketolation genes across animals (28, 40, 41). In birds, its expression influences the efficiency of the CYP2J19-BDH1L enzymatic complex. It may organize local membrane domains to optimize CYP2J19 functioning or facilitate substrate and cofactor delivery (above). TTC39B is not linked to vitamin A metabolism and does not interact with BCO1 or retinoid-binding proteins; its effect appears to be limited to the ketocarotenoid-producing cells (Fig. 1b; below): experimental knockdowns or natural variants of TTC39B reduce carotenoid output without affecting retinol levels (28, 40).

##### TRANSPORT AND STORAGE GROUP (FABPS, RBPS, CRABPS, PLINS)

Carotenoids are hydrophobic, requiring specialized carriers for transport and storage in aqueous environments. Several families of binding proteins perform these tasks in birds. The fatty acid-binding proteins (FABPs) are small intracellular proteins that chaperone carotenoids within cells (46, 47). The retinol-binding proteins (RBPs) include both intracellular forms that bind retinol/retinal, and the plasma RBPs (not studied here) that deliver retinol through the bloodstream (Supplementary Table S1) (14, 48). The cellular retinoic acid-binding proteins (CRABP1 and CRABP2) bind RA inside cells, regulating its availability to nuclear receptors or enzymes (49). The perilipins (PLINs) coat lipid droplets to manage carotenoid storage, long-distance transportation, and enzymatic access (50, 51).

###### FABPs

An evolutionarily conserved (46, 47) multigene family of proteins that bind hydrophobic ligands (Supplementary Table S1). Different FABPs are expressed in different tissues (Supplementary Table S1) and facilitate the intracellular transport of carotenoids (46, 52). For example, as intestinal enterocytes absorb carotenoids and some are cleaved to retinal by BCO1, FABPs may ferry these retinoids to the ER for esterification into vitamin A esters or into chylomicrons (53). Carotenoids that are incorporated into portomicron for secretion are transported by FABP1 and FABP2 by shuttling lipids to microsomal triglyceride transfer protein for portomicron assembly (47). In the liver, FABP1 can bind carotenoid metabolites and direct them toward Transport or Conversion groups (Fig. 1b). Some FABPs (e.g., FABP5) indirectly interact with nuclear receptors. For example, FABP5 binds RA and delivers it to PPARβ/δ in mammals (see Integration group, below), while CRABP2 delivers RA to RAR – creating a partition between proliferative (PPAR-mediated) and differentiative (RAR-mediated) RA signals (54, 55). PPARγ activation can upregulate FABP4 to enhance carotenoid storage in adipocytes (56, 57). Conversely, retinoic acid via RAR can counteract PPAR-induced effects; e.g., RA/RARα can suppress PPARγ target genes such as CD36 and FABP4 in liver to reduce lipid uptake (58, 59) (Fig. 1b). Thus, FABPs act at the intersection of metabolic and regulatory pathways (Fig. 1b) (55, 60).

###### RBPs

The RBP family includes both intra-and extracellular retinol carriers. Intracellularly, RBP1 and RBP2 are the main cytosolic retinol/retinal-binding proteins in vitamin A target tissues. RBP1 is widespread, binding retinol and retinal to facilitate either storage or oxidation to RA, while RBP2 is abundant in the small intestine, where it binds retinaldehyde produced by BCO1 and transports it for conversion to retinol and esterification (14, 27, 61). These binding proteins sequester retinol/retinal, preventing uncontrolled interactions and presenting them to different sets of enzymes either for storage or for RA production (14, 26). For example, in the avian liver, RBP1 ensures that vitamin A derived from carotenoids is esterified and stored in stellate cells or exported. In pigment-producing tissues, RBPs are typically low since retinoid production is minimized there (28). The cellular RBPs (RBP1/2/5/7; Supplementary Table S1) are closely linked to BCO1 expression (Fig. 1b). As mentioned above, in intestinal cells carotenoid cleavage by BCO1 is coupled with RBP2 binding of the retinaldehyde product serving as a prime example of an interaction between the conversion and transport groups (Fig. 1b): BCO1’s output immediately becomes the RBP2 cargo (26, 27). The presence of RBP in BCO1-expressing cells is so consistent that these proteins are jointly regulated in vitamin A processing tissues (48). On the other hand, RBPs do not usually occur in cells with ketocarotenoid production (e.g., feather follicles expressing CYP2J19) (28), further highlighting the separation of the two conversion group pathways (Fig. 1b).

###### CRABPs

CRABP1 and CRABP2 are short (Supplementary Data S1), evolutionarily conserved cytosolic proteins that bind RA. Their role is to sequester RA and regulate its access to nuclear receptors (RAR/RXR) and to enzymes (CYP26 family) (49). CRABP1 is ubiquitously expressed, while CRABP2 is commonly found in tissues with active RA signaling (e.g., integument). Birds likely use CRABPs to modulate retinoic acid derived from carotenoids: CRABP2 delivers RA to RAR in the nucleus, enhancing RA-mediated gene transcription, CRABP1 sequesters RA in the cytosol or facilitates its degradation, thereby decreasing RAR activation (26, 27, 49). CRABPs thus act as intracellular switches that influence functional consequences of carotenoid-derived RA directing RA towards different fates – towards gene activation and towards catabolism. CRABP1 in the same cell might ensure that any excess RA is degraded to prevent overactivation of RA signaling (49). CRABPs link directly with the Integration group (Fig. 1b) by modulating how much RA reaches RAR/RXR. Specifically, CRABP2 enhances RAR signaling by presenting RA to the receptor (49), whereas CRABP1 can limit RA delivery to RAR; FABP5, in turn, can direct RA toward PPARβ/δ signaling. The interaction of CRABPs and FABP5 is an example of network-wide integration enabled by the transport group proteins (Fig. 1b): depending on which binding protein dominates, RA can activate different nuclear pathways (RAR vs PPAR), leading to different gene expression patterns. For carotenoid metabolism feedback, CRABP2 promotes the RAR-mediated induction of ISX, thereby linking high carotenoid intake to downregulation of dietary absorption and enzyme genes (Fig. 1) (15, 16).

###### PLINs

Perilipins coat intracellular lipid droplets, thereby stabilizing carotenoid storage and regulating access to stored carotenoids for metabolism, transportation and mobilization (50, 62, 63). Here we studied PLIN1, PLIN2 and PLIN3 which are abundant in many tissues (Supplementary Table S1). In avian carotenogenesis, in addition to storing large amounts of carotenoids for later utilization (e.g., during oogenesis or molt), perilipins also mitigate cell membrane oxidative damage by sequestering free carotenoids in droplets. PLIN2 is particularly well known to associate with carotenoid-rich droplets in liver and skin and also participates in carotenoid storage in other vertebrates (51, 64). PLINs sequester and isolate carotenoid metabolites of various metabolic derivations from enzymes in the Conversion group (Fig. 1b) (51, 65, 66). This physical separation also allows birds to accumulate large amounts of excess carotenoids without converting or breaking them down. Hormonal signals trigger lipase access to these droplets, at which point carotenoids can be released and transported out to the Conversion or other functional groups. For example, PLIN1 in adipocytes prevents basal lipolysis, but allows lipases to act upon phosphorylation (50). In adipose tissues, starvation can trigger mobilization of fat, which also releases carotenoids, that can affect subsequent pigmentation (51). Further, PPARγ (Integration group, below) can upregulate PLINs, promoting lipid storage (67): activation of PPARγ facilitates carotenoid storage by inducing droplet formation and perilipin coating, whereas activation of RAR (by RA) might reduce fat storage and associated carotenoid storage. Thus, PLIN proteins are fundamental to the utilization vs. storage decision: once carotenoids are enclosed in a PLIN-coated droplet, they are temporarily removed from circulation and enzymatic conversion (51) (Fig. 1b).

##### INTEGRATION GROUP (NUCLEAR REGULATION: PPARγ, RAR α/β/γ, ROR α/β/γ, ISX)

These nuclear factors integrate signals from carotenoid uptake, conversion, allocation and transport to regulate gene expression and maintain carotenogenesis (Supplementary Table S1, Fig. 1b).

###### PPARγ

A ligand-activated transcription factor that forms a heterodimer with RXR. It is a master regulator of adipogenesis and lipid metabolism, controlling genes involved in fatty acid uptake, triglyceride storage, and glucose metabolism (57). PPARγ’s ligands include fatty acids; through RXR, it can also be influenced by vitamin A. Thus, carotenoid derivatives modulate PPARγ indirectly: when abundant vitamin A leads to high 9-cis RA, RXR is activated and partners with PPARγ, enhancing its expression (68). Conversely, all-trans RA via RAR can antagonize PPARγ’s effects by competing for RXR or inducing factors that inhibit PPAR pathways (56, 60). When carotenoid supply is high, PPARγ is active and promotes adipose tissue formation, creating greater storage capacity for lipids and carotenoids; for example, it upregulates CD36 and promotes downstream lipid droplet accumulation regulated by FABP4 (Fig. 1b) (60). As mentioned above, PPARγ also induces PLIN1 expression to form and stabilize new lipid droplets (Fig. 1b) (67). The net effect is that PPARγ activation channels carotenoids into storage in relation to uptake and concentration (Fig. 1) (57). PPARγ closely interacts with RARs (Fig. 1b): under vitamin A-rich conditions, RA (via RAR) can suppress PPARγ activity; activating RARα in the liver prevents PPARγ from upregulating CD36, thereby reducing carotenoid accumulation (58, 59). Both PPARγ and RARs share RXR as a heterodimer partner, so an excess of RA can drive RXR to pair with RAR instead of PPARγ, diminishing PPARγ’s transcriptional activity (56). PPARγ and RAR have opposing target genes: RAR (through ISX) downregulates SCARB1 and BCO1, whereas PPARγ upregulates CD36 and FABP4 to integrate uptake, utilization and storage (Fig. 1b). PPARγ also indirectly promotes retinoid availability; activation of PPARγ in some cells induces RBP7 (see Transport group) that can serve as a PPARγ cofactor (26, 27, 61).

###### RARs (α/β/γ)

The RARs are nuclear receptors that bind RA and regulate gene expression by binding RAR response elements (RAREs) in DNA as RAR/RXR heterodimers (69, 70). In the context of carotenoid metabolism, RARs serve as the sensors for vitamin A. When dietary carotenoids are abundant and more RA is produced, RARs are activated and trigger feedback mechanisms (as noted above) (15, 16). RARα helps control genes involved in lipid metabolism: activating RARα can suppress PPARγ and its targets (e.g., CD36), which affects how much carotenoid-rich lipids are taken up by the liver and the gut (58, 59). RARs are closely functionally integrated with RXR. Together, these receptors can induce transcriptional repressors or activators that impact ISX and genes for RA degradation and some binding proteins like CRABP2 (49). In the developing eye, RA signaling – transduced via nuclear receptors such as RARβ, RARγ, and RXRs – regulates ocular morphogenesis (71) and photoreceptor subtype balance (72), whereas cone oil droplet biogenesis relies on coordinated neutral lipid and carotenoid metabolism within maturing cones (73). Similarly, knockouts of RARα or RXRα reproduce features of vitamin A deficiency in mice (74, 75).

###### RORs (α/β/γ)

RORs are orphan nuclear receptors structurally related to RARs. In vertebrates, RORβ regulates photoreceptor development (76) integrating carotenoid accumulation in cones’ oil droplets (77). Overall, RORs indirectly integrate carotenoid utilization with developmental and hormonal regulation.

###### ISX

A transcription factor that is the molecular bridge between the Absorption and Conversion functional groups by acting as a gatekeeper for intestinal carotenoid absorption (Fig. 1b). It is expressed in the small intestine (Supplementary Table S1) and is directly regulated by RA (above) (16). Once produced, ISX protein binds to the promoters of SCARB1 and BCO1 and represses their transcription (15, 16). RAR–RXR heterodimers induce ISX, which suppresses SCARB1 and BCO1 (Absorption and Conversion groups). This feedback is crucial for diet-dependent carotenoids with unpredictable availability; it both prevents vitamin A toxicity and may ensure that some carotenoids remain unconverted and channeled to the pigmentation pathway when vitamin A is sufficient (15, 16). ISX is the central target of a regulatory network discussed above (Fig. 1b).

### Supplementary Text S2

#### Robustness tests of the homotypic ε_ST_ metric

FINCHES is a mean-field model parameterized for intrinsically disordered regions (IDRs) (78), and ε_ST_ is the homotypic interaction parameter ε divided by sequence length (Methods). The fatty acid-binding proteins (FABPs), the cellular retinol-binding proteins (RBPs) and the cellular retinoic acid-binding proteins (CRABPs) in the transport layer are short (123-142 residues), compact β-barrel proteins (Supplementary Text S1, Supplementary Data S1). We therefore tested (i) whether the FINCHES assessment of binding avoidance is biased by protein length or net charge; (ii) whether the differences among layers persist when ε is computed only on disordered or only on solvent-exposed residues; and (iii) whether ε_ST_ differs from what amino-acid composition alone predicts.

##### (i) ε_ST_ does not depend on protein length or net charge

FINCHES computes ε for a homotypic pair as ε = L × (W̅ − 2b), where b is the baseline term and the weighted pairwise interaction matrix W̅ is the mean of W, such that ε_ST_ = ε/L = W̅ − 2b is the mean pairwise interaction parameter of the sequence. Tandem duplication of each of the 3,534 unique sequences, which doubles L at constant composition, changed ε_ST_ by 0.07 × 10^−3^ on average (max 1.4) vs e.g., a difference of 36 × 10^−3^ between the transport and specialized layers. When the 20 proteins of at least 264 residues were cut into contiguous 131-residue windows (the median length of within-cell carriers; 1,771 species × protein sequences), the mean ε_ST_ of the windows fully reproduced the full-length value (*r* = 0.95; mean shift 2.9; Supplementary Fig. S6d). Windows of absorption, conversion and distribution layer proteins averaged -5.4 (full length: -5.9), and 0.1% of them reached the level of FABP1, FABP4 and FABP5 (≥ 45), compared with 46-67% of PLIN windows. Thus, short length of these proteins does not result in high ε_ST_.

Across all proteins, ε_ST_ nevertheless decreases with length (*r* = -0.35 with log_10_ length, *P* = 0.051; Spearman ρ = -0.50, *P* = 0.004; Supplementary Fig. S6a, Supplementary Table S2). The same tendency is shown by the proteins of individual species (median ρ = -0.40). This association, however, reflects the composition of proteins of different size. First, at nearly constant length (123-142 residues) and with the same fold, the 11 within-cell carriers span most of the range of all other proteins: from -1.2 to 53.9. Second, in a regression of ε_ST_ on log_10_ length, NCPR, FCR and membership of the transport layer, transport membership was retained (*β* = 33.0 ± 9.1, *P* = 0.001; R^2^ increased from 0.17 to 0.45) but length was not (*P* = 0.16).

ε_ST_ was unrelated to net charge per residue (NCPR; *r* = 0.07, *P* = 0.73; Supplementary Fig. S6b; or its absolute value *r* = 0.06, *P* = 0.76), related to the fraction of charged residues (FCR; *r* = 0.36, *P* = 0.049) and strongly and negatively related to the fraction of aromatic residues (*r* = -0.71, *P* <0.001; Supplementary Fig. S6c). Switching off the electrostatic term of the force field (Methods) largely preserved the ranking of proteins (*r* = 0.98) and ε_ST_ increased in every layer (by 5-13 × 10^−3^): in these near-neutral sequences, attraction between opposite charges slightly outweighs repulsion between like charges. The self-avoidance of carriers is therefore not an effect of net charge. In this force field it marks sequences that are poor in aromatic residues: 3.8-5.8% F + W + Y in PLINs and 6.1-7.6% in FABP1, FABP4 and FABP5, against 8.9-12.7% in absorption and conversion layer proteins.

##### (ii) ε_ST_ of disordered, folded and solvent-exposed residues

Using the per-residue MetaPredict scores, we then recomputed ε_ST_ over IDR residues and over folded residues separately. The weighted matrix W was built on the full chain, so that the sequence-context weights were retained, and then averaged over the respective block of residues; at least 20 residues were required. IDR-only values could be estimated for 17 proteins (7 to 154; Supplementary Fig. S7a, Supplementary Table S2). IDRs are thus self-avoiding in all layers, including the short terminal IDRs of absorption and distribution proteins (layer means 71 ± 14 and 154 ± 6; Supplementary Table S3), and the IDRs of PLINs (55 ± 4) are not more self-avoiding than these (PLINs vs absorption = -14 ± 8, *P* = 0.094; PLINs vs distribution = -92 ± 23, *P* < 0.001). Across proteins, IDR-only values were uncorrelated with full-length values (*r* = 0.02). Within-cell carriers have almost no IDRs (0-6% of residues). Folded-only values, in contrast, reproduced the full-length pattern (*r* = 0.87 across 31 proteins): PLINs 32.6 ± 1.5 and within-cell carriers 14.9 ± 5.5, against absorption -7.3 ± 0.9, conversion -11.8 ± 1.1 and distribution -5.1 ± 1.0 (transport vs absorption = 35.8 ± 0.6; transport vs conversion = 41.5 ± 4.4; both *P*s < 0.001). The difference between carriers and specialized layers reported in Figs. 2 and 3 therefore is not biased by relative IDR span. This raises the question of whether it is an artifact of applying FINCHES to folded domains, and of which folded residues are available for intermolecular contact.

Solvent exposure was taken from experimental structures: for each of the 11 within-cell carriers, the crystal or NMR structure of a mammalian orthologue (Supplementary Table S4). DSSP accessibilities were converted to relative solvent accessibility (RSA) and transferred to each avian sequence by pairwise alignment (59-93% identity for orthologous templates, 36-43% for paralogous templates). ε_ST_ was then recomputed over exposed residues (RSA ≥ 25%) in two ways: with the sequence-context weights of the full chain, or with the FINCHES convention for folded domains, in which surface residues were treated as isolated beads. Two independent crystal structures of FABP4 gave the same exposure class for 94% of residues. Restricting ε to exposed residues made most within-cell carrier more self-avoiding (Supplementary Fig. S7b, Supplementary Table S4). ε_ST_ was positive for all 11 proteins (nine of 11 on full sequences), and the layer mean rose from 14.5 ± 5.6 to 54.9 ± 9.2 (35.7 ± 7.7 with isolated beads; 82 species), whereas buried residues were self-attracting (-41.0 ± 9.4). Exposed residues of these proteins contain 1-4% aromatic residues, buried residues 12-23%. The self-avoidance of FABPs, CRABPs and RBPs is thus a property of their solvent-exposed surface and not an artifact of treating a β-barrel as a disordered chain.

The same restriction also raised ε_ST_ of the folded proteins of the specialized layers: CD36 from 2.6 to 47.1 (29.1 with isolated beads), SCARB1 from -7.0 to 25.2 (18.9) and BCO1 from -3.7 to 50.5 (21.4); only BCO2 remained self-attracting (-21.4 to -8.6; -16.6). At the layer level, absorption (CD36 and SCARB1) changed from -3.4 ± 2.0 to 34.3 ± 3.0 (22.2 ± 2.9) and conversion (BCO1 and BCO2; 14 species) from -11.1 ± 1.4 to 25.4 ± 3.5 (5.3 ± 2.5). Thus, negative full-sequence ε_ST_ of the specialized layers reflects their aromatic-rich buried cores and the ordering of layers is replicated. Within-cell carriers remained more self-avoiding than the conversion enzymes at every threshold and under both conventions (difference at RSA ≥ 25%: 35.2 ± 3.9, *P* < 0.001 with sequence context and 38.0 ± 2.8, *P* <0.001 with isolated beads; full sequences: 31.4 ± 2.1, *P* <0.001). Against absorption proteins, the difference on exposed residues was of the same size as on full sequences but less certain at the 25% threshold (full sequences: 17.2 ± 6.2, *P* = 0.007; exposed: 19.6 ± 9.9, *P* = 0.051 with sequence context and 12.2 ± 8.0, *P* = 0.13 with isolated beads), and it was larger and significant at RSA ≥ 10% (32 and 27; both *Ps* <0.001) and at RSA ≥ 40% (65 and 60; *P* = 0.002 and *P* = 0.003). On solvent-exposed surfaces, carriers are therefore more self-avoiding than the conversion proteins and, at the 10% and 40% thresholds, than the absorption proteins.

##### (iii) Composition-preserving shuffled-sequence null

Each of the 3,534 unique sequences was shuffled (*n* = 500), preserving length and amino-acid composition, and ε_ST_ was recomputed. In FINCHES, sequence order affects ε only through the local weighting of clustered charges and clustered aliphatic residues, so this null specifically isolates the contribution of composition. The observed ε_ST_ of the 31 proteins was fully replicated (*r* = 0.986, Supplementary Fig. S8, Supplementary Table S2). The mean absolute difference between observed and null values was 2.4 × 10^−3^, against a s.d. of 19.4 among proteins, so that composition accounts for 97% of the variance among proteins. It also reproduces the contrasts among layers (Supplementary Table S3): for example, transport vs absorption = 34.7 under the null against 36.5 observed, and transport vs conversion = 43.3 against 46.0. Sequence order adds a small consistent increment in the three PLINs (observed values above the central 95% of shuffles in 88-100% of sequences; +4 to +7 × 10^−3^; mean z = 2.7-3.5) and a small decrease in CRABP2 (-8.3; z = -2.4). All other transport proteins, including all FABPs, remain within their null distributions. The differences among layers are thus robust to sequence order: they do not depend on particular motifs, isoforms or annotation of termini (Supplementary Figs. S6c and S8).

#### Methods for tests in this section

We used 2,734 species × protein entries in Supplementary Data S3. Sequences with non-standard residues were excluded; values were averaged over all isoform records of a protein within a species, as in Supplementary Data S1. ε was computed with the Mpipi-GG parameters and default charge and aliphatic weighting of FINCHES. NCPR and FCR count K and R as +1 and D and E as -1. For the electrostatics-off calculation the pair parameters were recomputed from the Wang–Frenkel term. For windows, the longest isoform of each species × protein entry of ≥ 264 residues was scanned with 131-residue windows in steps of 11 residues. Intrinsic disorder was predicted with the MetaPredict V3 network. Solvent accessibilities were taken from the DSSP databank (79, 80) of PDB-REDO for the first chain of each template (Supplementary Table S4) and divided by the theoretical maximum accessibility of each residue type (81). Templates were aligned to avian sequences with the Needleman–Wunsch algorithm (BLOSUM62; gap opening -10, extension -0.5; free end gaps), and only aligned positions were used. Layer values, λ-optimized GLS means, phylogenetic paired contrasts, phylogenetic transformation, Lindeman-Merenda-Gold decomposition were calculated as described in the Methods of the main text.

**Supplementary Figure S1.**
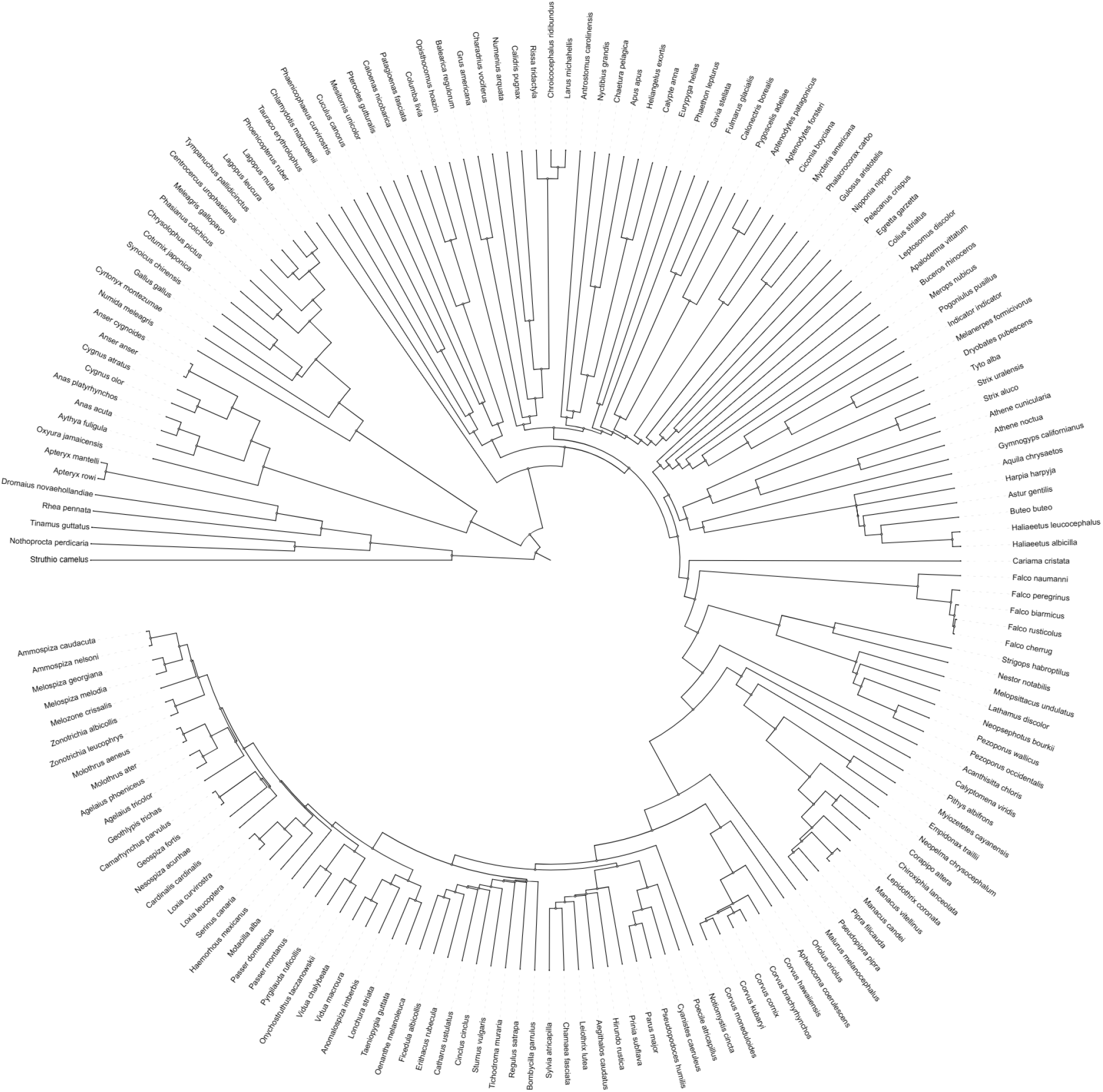
Phylogeny of 172 species under this study (Supplementary Data S2)

**Supplementary Figure S2.**
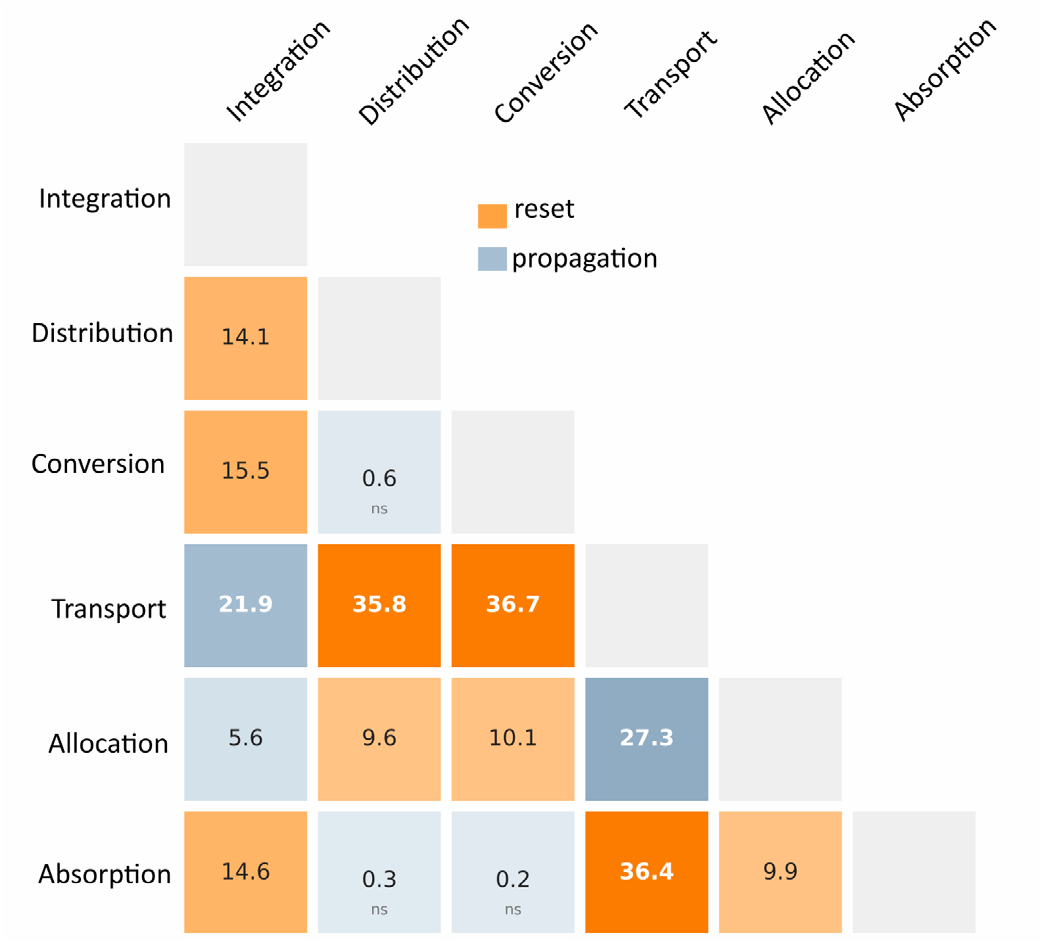
Cumulative magnitude of resets |Δ| ×10³ between functional groups calculated from phylogenetic linear contrasts. ns – multiple-range *p* > 0.05.

**Supplementary Figure S3.**
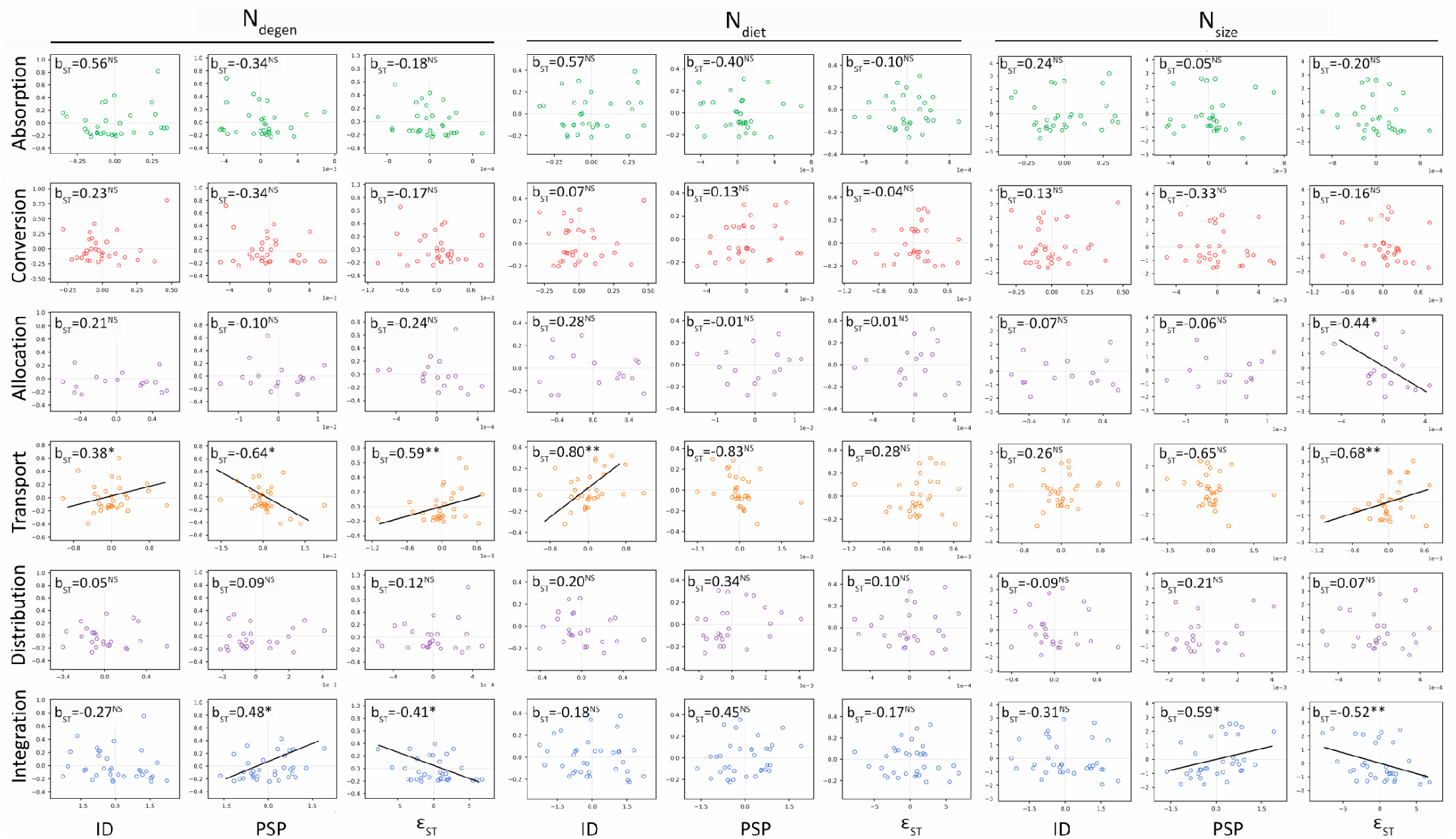
Partial regression plots of phylogenetically transformed biophysical properties of proteins (ID, PSP, ε_ST_) in six functional groups against number of degenerate nodes (Ndegen), number of dietary nodes (Ndiet) and network overall size (PC1 of nodes and edges). b_ST_ is standardized regression coefficient, \**p* < 0.1, \*\**p* < 0.05.

**Supplementary Figure S4.**
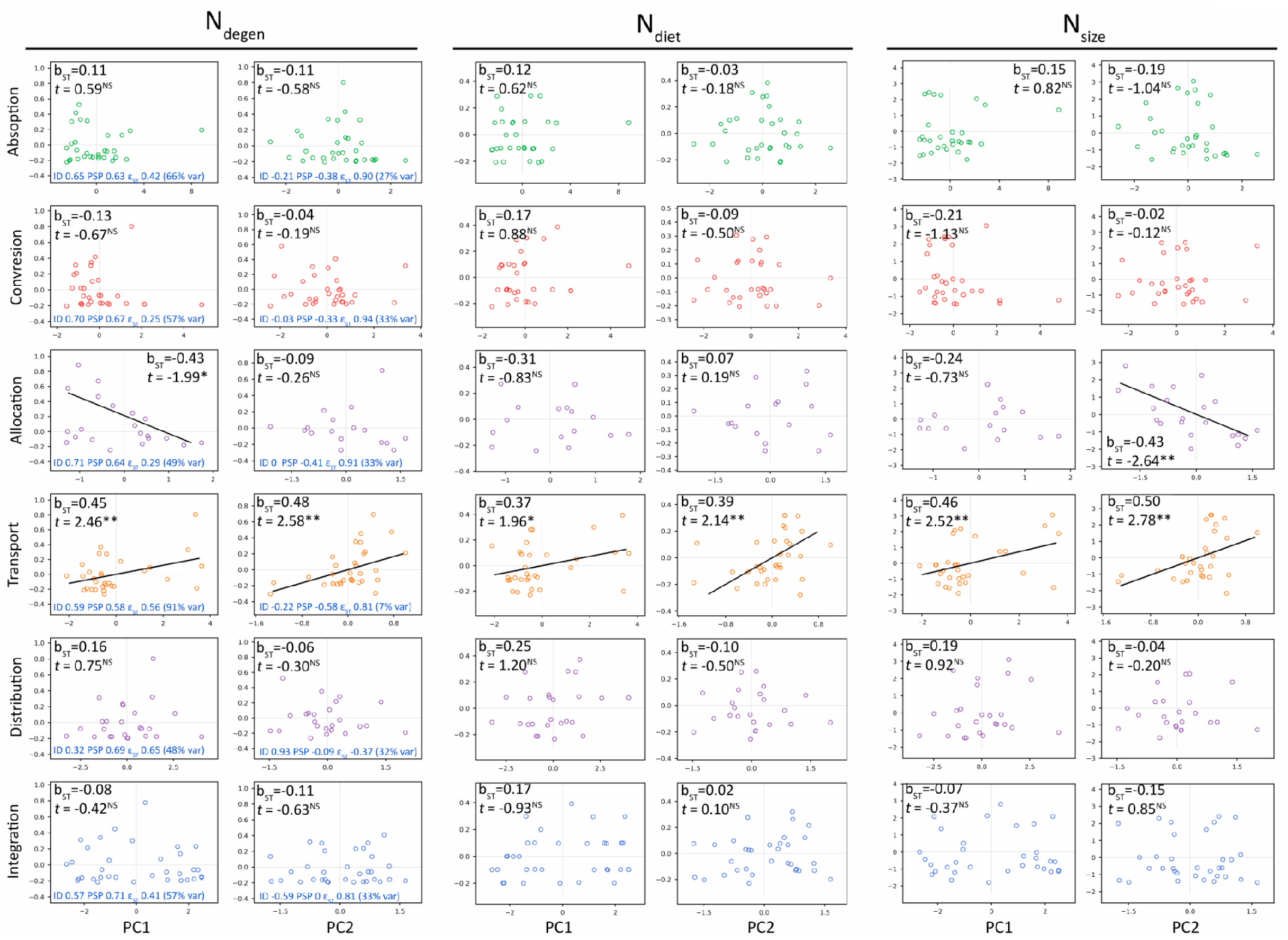
Partial regression plots of *within functional group* principal components of phylogenetically transformed biophysical properties of proteins against number of degenerate nodes (Ndegen), number of dietary nodes (Ndiet) and network overall size (PC1 of nodes and edges). b_ST_ is standardized regression coefficient, \**p* < 0.1, \*\**p* < 0.05. Numbers in blue in Ndegen panels show principal decomposition of biophysical properties for the group and percent of explained variance. Except for Distribution group, PC1 is a general disorder and PC2 is ε_ST_ in excess of that explained by disorder.

**Supplementary Figure S5.**
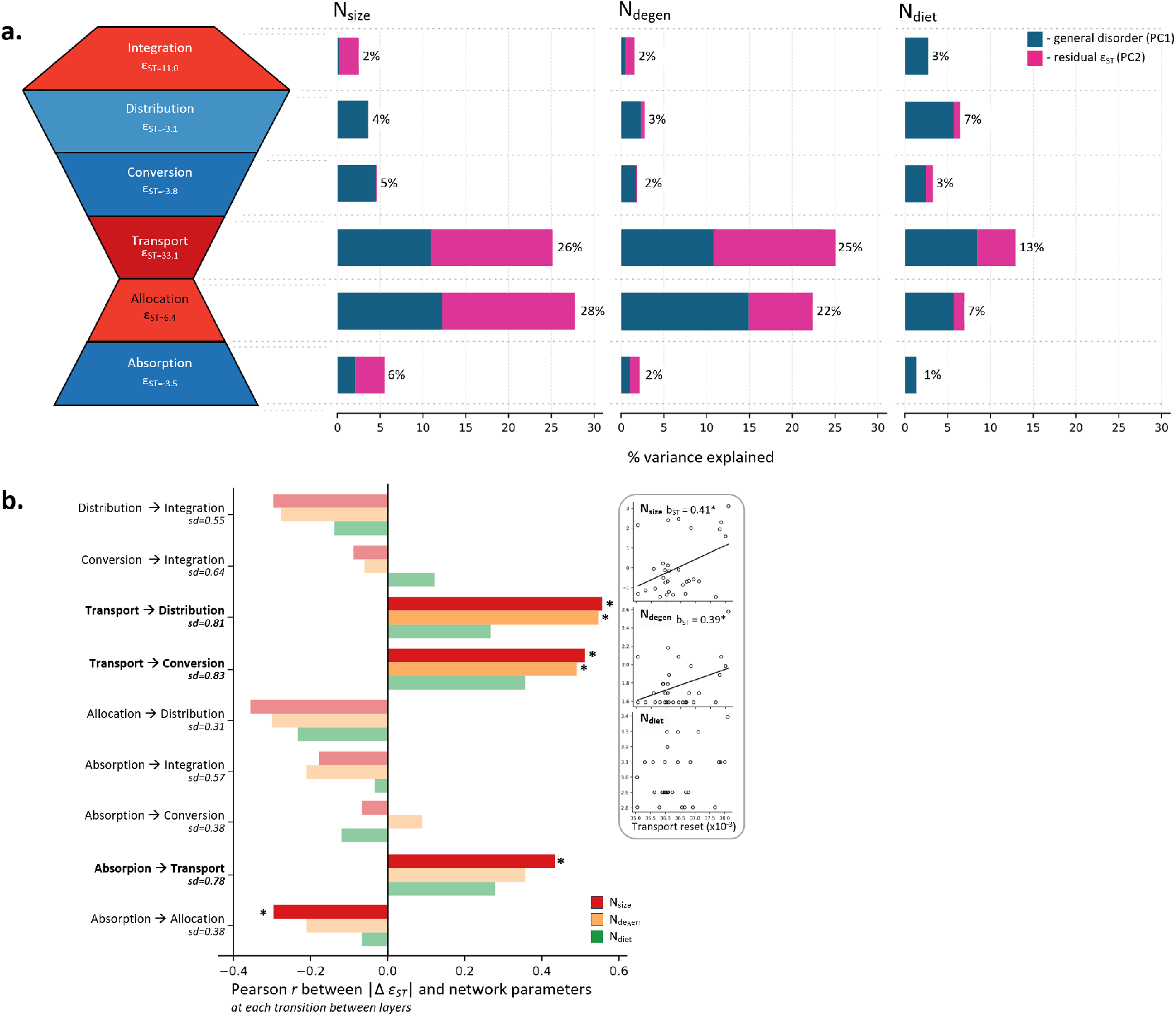
Binding self-avoidance of the waist layers drives network growth and connectivity by resetting specialization and inducing pathway branching. (a) Percentage of among-species variance in network size (Nsize), number of degenerate nodes (Ndegen) and number of dietary carotenoids (Ndiet) explained by within-group principal components of protein biophysical properties (Supplementary Fig. S4 shows the decomposition); stacked bars are partial R^2^ (Methods) aligned to the bow-tie layers at left (layer colors and ε_ST_ as in Fig. 3a). (b) Resets in binding regimes between functional complexes that contribute to network changes. Insets show partial regression plots of network parameters on the magnitude of the reset, |Δε_ST_| at the transport layer.

**Supplementary Figure S6.**
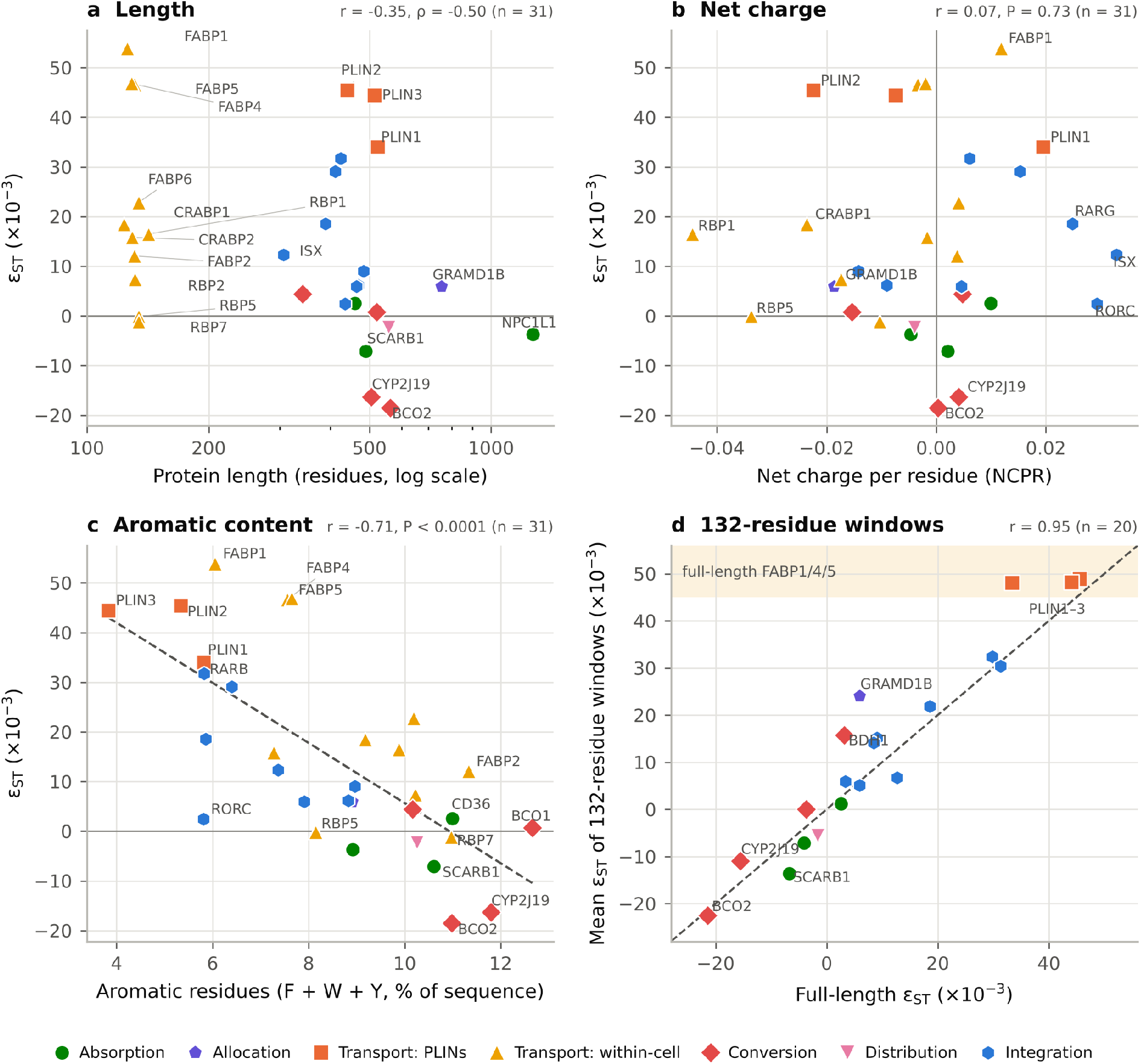
ε_ST_ in relation to protein length, net charge and aromatic content. Points are protein means across species (Supplementary Data S1); colors and symbols denote functional layers. (a) ε_ST_ against protein length. (b) ε_ST_ against net charge per residue. (c) ε_ST_ against the percentage of aromatic residues, with the least-squares line. (d) Mean ε_ST_ of contiguous 132-residue windows against the full-length value for the 20 proteins of ≥ 264 residues; dashed line, 1:1; shaded band, range of full-length FABP1, FABP4 and FABP5. NCPR, aromatic content and windows are computed from Supplementary Data S3.

**Supplementary Figure S7.**
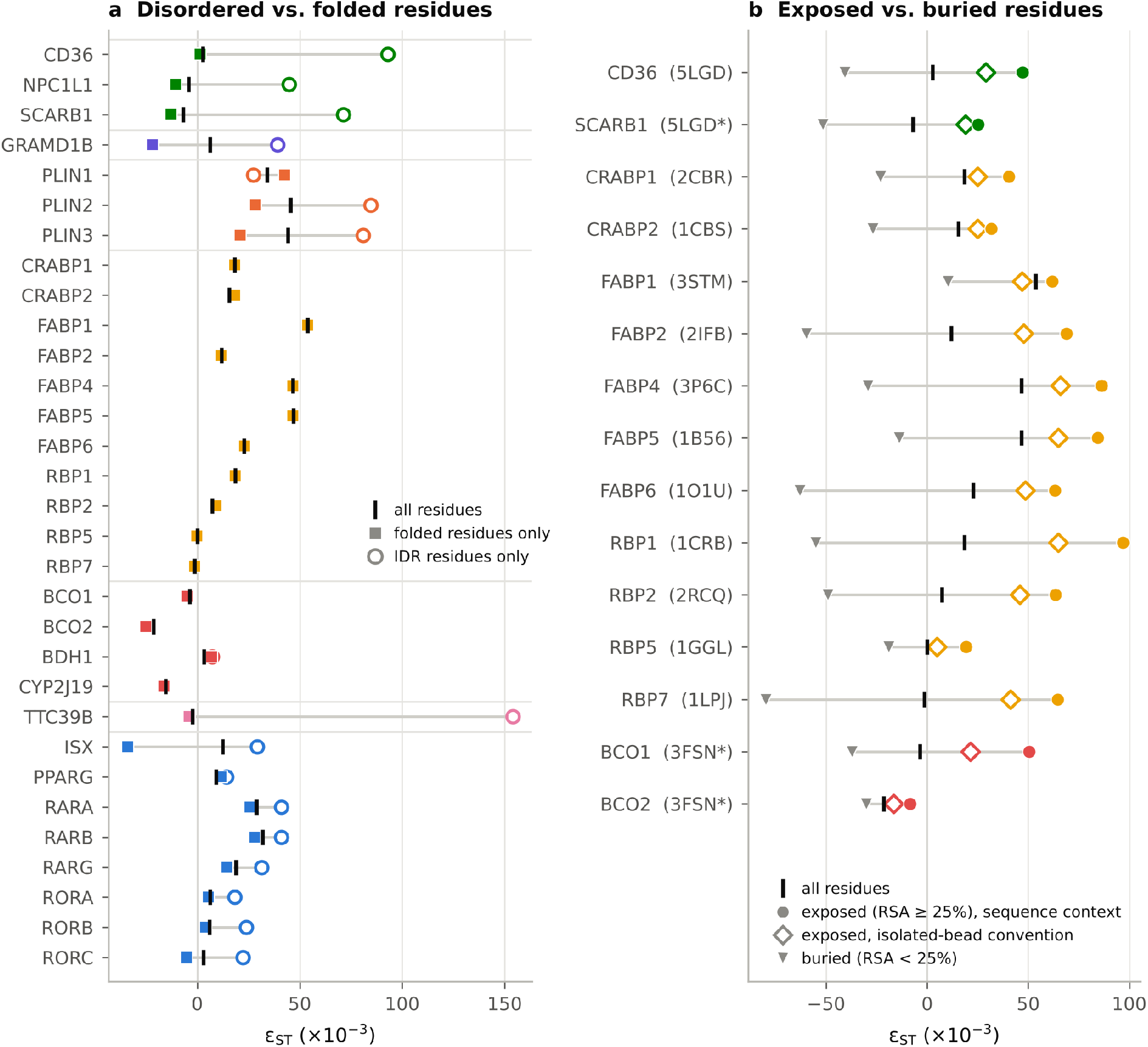
ε_ST_ of residue subsets. **(a)** ε_ST_ of all residues (black bar), folded residues only (filled squares) and IDR residues only (open circles) for the 31 proteins; values are means across species, colors denote functional layers. **(b)** ε_ST_ of solvent-exposed residues (RSA ≥ 25%) computed with sequence-context weights (filled circles) or with surface residues treated as isolated beads (open diamonds), and of buried residues (grey triangles), for the 15 proteins with an experimental structural template (PDB code in parentheses; asterisks mark paralogous templates; Supplementary Table S4).

**Supplementary Figure S8.**
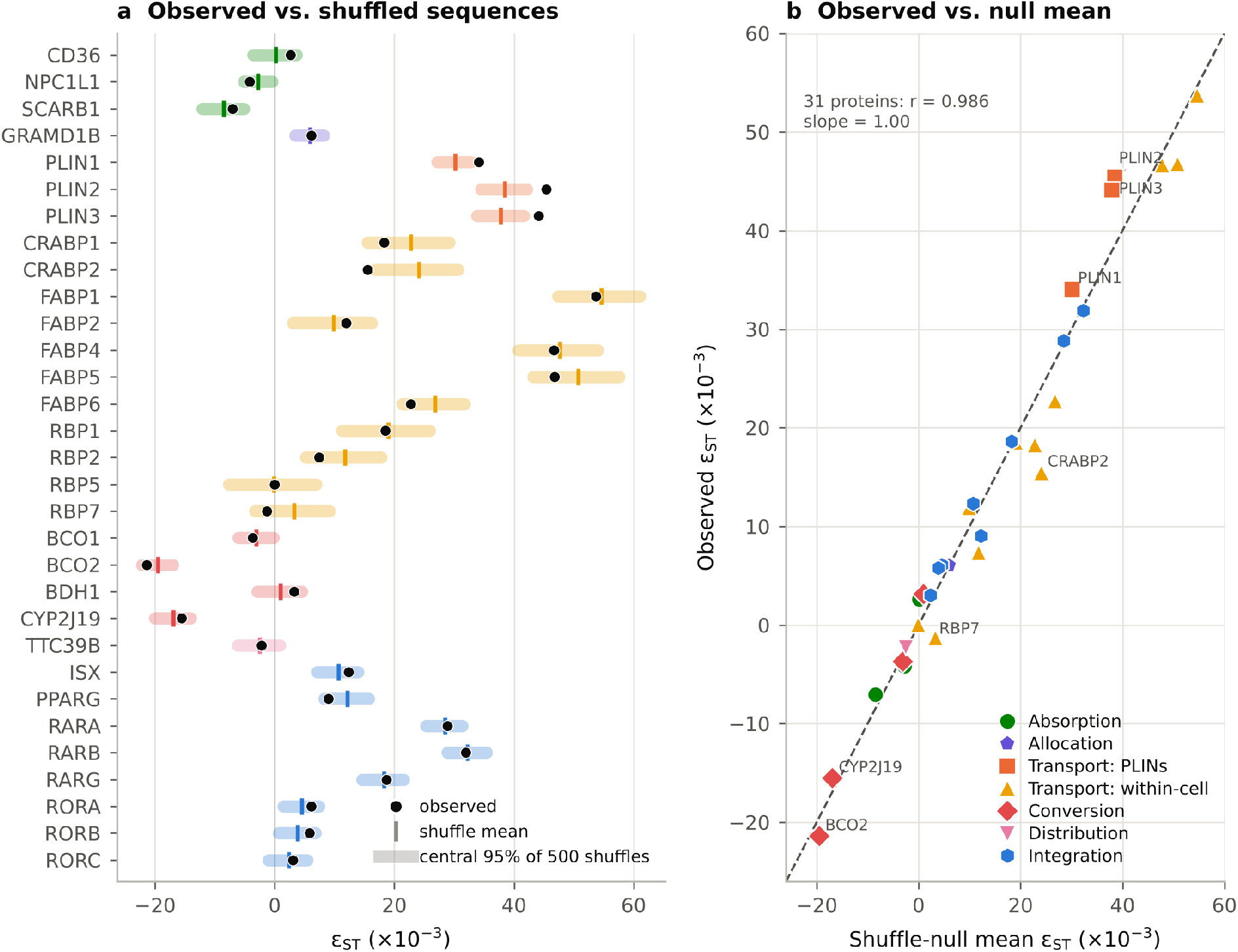
Observed ε_ST_ compared with a composition-preserving null. **(a)** Observed ε_ST_ (black points), mean of 500 shuffles (colored bars) and 95% of shuffles (shaded) for each protein; values are means across species. **(b)** Observed ε_ST_ vs the mean of the shuffle null; dashed line, 1:1. Based on Supplementary Data S3.

**Supplementary Table S1.** Main function and expression sites for core avian carotenogenesis proteins. Functional group designation follows Fig. 1b based on Supplementary Text S1.

| Group | Protein/Gene<br>(species with full sequence) | Main function | Key expression sites | Key Refs. |
|---|---|---|---|---|
| **Dietary absorption** | **CD36**<br>(FAT/CD36)<br>*n* = 164spp | Scavenger receptor aiding uptake of carotenoid-containing micelles and lipoproteins. Absorption and peripheral tissue uptake in mammals. | Intestinal enterocytes (microvillar membrane); peripheral tissues (e.g., liver, adipose, retina, macrophages). | (5, 18) |
|  | **SCARB1**<br>(SR-BI)<br>*n* = 167spp | Membrane receptor for selective uptake of dietary carotenoids from micelles; intestinal absorption in birds. | Intestinal brush-border enterocytes; liver hepatocytes and cell membranes, retinal cells. | (8, 9, 11) |
|  | **NPC1L1**<br>*n* = 96spp | Intestinal carotenoid uptake via vesicular endocytosis and cholesterol transporter, particularly in mammals. | Apical brush-border membrane of intestinal enterocytes; liver. | (7, 19) |
| **Allocation** | **GRAMD1B**<br>(Aster-B)<br>*n* = 149spp | Lipid-transfer protein that shuttles carotenoids at membrane contact sites. Directs absorbed carotenoids to modification sites. | ER–plasma membrane sites across many cell types and tissues. | (37, 44, 82) |
| **Distribution** | **TTC39B**<br>*n* = 162spp | Enables or enhances ketocarotenoid production; co-factor of regulatory proteins in pigment cells. | Co-expressed with “conversion group” (e.g., CYP2J19 in pigment-producing tissues). | (28, 40) |
| **Chemical conversion** | **BCO1**<br>(β-Carotene 15,15′-dioxygenase)<br>*n* = 166spp | Enzyme that symmetrically cleaves provitamin A carotenoids to yield retinal. Provides vitamin A for vision and systemic use. | Cytosol of intestinal cells (for dietary β-carotene cleavage); also liver, retina, and other tissues requiring retinaldehyde production. | (12, 83) |
|  | **BCO2**<br>(β-Carotene 9′,10′-oxygenase)<br>*n* = 155spp | Enzyme that asymmetrically cleaves carotenoids at alternative bonds. Initiates carotenoid degradation, disposes excess carotenoids, and prevents over-accumulation. | Mitochondrial inner membrane of cells of tissues that limit carotenoid accumulation (e.g., liver, retina), low in pigmented tissues. | (23, 33) |
|  | **CYP2J19**<br>*n* = 6spp | Cytochrome P450 ketolase with broad catalytic capacity for carotenoid conversion. | ER membranes of cells in carotenoid-depositing tissues (feather follicles, red oil droplets in retinal cone cells). | (22, 29, 30) |
|  | **BDH1L**<br>(3-Hydroxybutyrate dehydrogenase-like)<br>*n* = 159spp | Dehydrogenase that oxidizes hydroxyl groups on carotenoid rings to ketones. Acts sequentially after CYP2J19. | Cytosol (or membrane-associated) in the same pigment-producing cells as CYP2J19. | (22, 28, 42, 84) |
| **Transport / Storage** | **FABPs**<br>(Fatty Acid-Binding Proteins)<br>*n* = 64spp | Intracellular lipid-binding proteins that can transport and store carotenoids to lipid droplets. | Cytosol of specific cells: FABP1– liver and small intestine; FABP2 –glandular stomach, the duodenum and ileum; FABP4 – adipose tissue; FABP5 – liver, epithelial tissues/storage; FABP6 – liver and the ileum. | (18, 46, 47) |
|  | **PLINs**<br>(Perilipins)<br>*n* = 169spp | Coat lipid droplets, stabilizing, regulating and storing carotenoids. Regulating lipase access to enclosed carotenoids. Transport and sequestration of carotenoids in cytosolic storage depots until mobilization. | Lipid droplets in adipose tissue, liver, and steroidogenic cells; PLIN1 in adipocytes, PLIN2 and PLIN3 in cells of tissues with active lipid storage. | (50, 51) |
|  | **RBPs**<br>(Retinol-Binding Proteins)<br>*n* = 77spp | Bind and transport vitamin A compounds: RBP1/2/5/7 bind retinal/retinol in cells (sequentially after BCO1), transporting retinol to metabolic pathways or to storage. | RBP1/5 – retinol storage in liver, eye, kidney; nervous system and limb buds in embryos; RBP2 – small intestine; RBP7 – adipose-specific | (14, 48, 85) |
|  | **CRABPs**<br>(Cellular Retinoic Acid-Binding Proteins)<br>*n* = 53spp | Bind all-trans retinoic acid (RA) in the cytosol, buffering RA levels and modulating its availability to nuclear receptors or catabolic enzymes. CRABP1 channels RA to degradation, CRABP2 – to RAR for gene regulation. | Cytosol (and nucleus) of many RA-responsive tissues (e.g. intestinal mucosa, liver, skin). Coexpressed with active RA metabolism. | (49, 55, 86) |
| **Integration/Nuclear regulation** | **PPARγ**<br>(Peroxisome proliferator-activated receptor gamma)<br>*n* = 16spp | Nutrient-sensing nuclear receptor, integrates genes involved in lipid uptake, adipose formation, and lipid metabolism. Links metabolic state to carotenoid absorption, storage, and transport. | Nucleus of adipocytes, macrophages, and other lipid metabolizing cells. | (57, 87) |
|  | **RARα/β/γ**<br>(Retinoic Acid Receptors)<br>*n* = 92spp | Nuclear receptors activated by RA. Regulate and integrate numerous carotenoid metabolism genes (feedback regulation of uptake and cleavage, e.g., in vitamin A homeostasis and developmental regulation of carotenoid use). | Nuclei of cells in numerous tissues (subtype- isoform-specific patterns) involved in feedback regulation of carotenogenesis. | (69, 70) |
|  | **RORα/β/γ**<br>(Retinoic acid receptor-related Orphan Receptors)<br>*n* = 168spp | Orphan nuclear receptors that indirectly regulate and integrate carotenoid allocation to vision vs other organismal functions via metabolic signaling pathways. | Nuclei of specific cell types: RORβ - retinal neurons (cones) and circadian centers; RORα - widespread (e.g. cerebellum, muscle), RORγ - in immune cells. | (68, 76, 88) |
|  | **ISX**<br>(Intestine-Specific Homeobox)<br>*n* = 135spp | Transcription factor acting as a feedback regulator of carotenoid absorption, allocation and chemical conversion in response to RA levels. | Nuclei of intestinal epithelium cells (primarily in the small intestine). Expression is induced by RA via RAR signaling. | (15, 16) |

**Supplementary Table S2.**
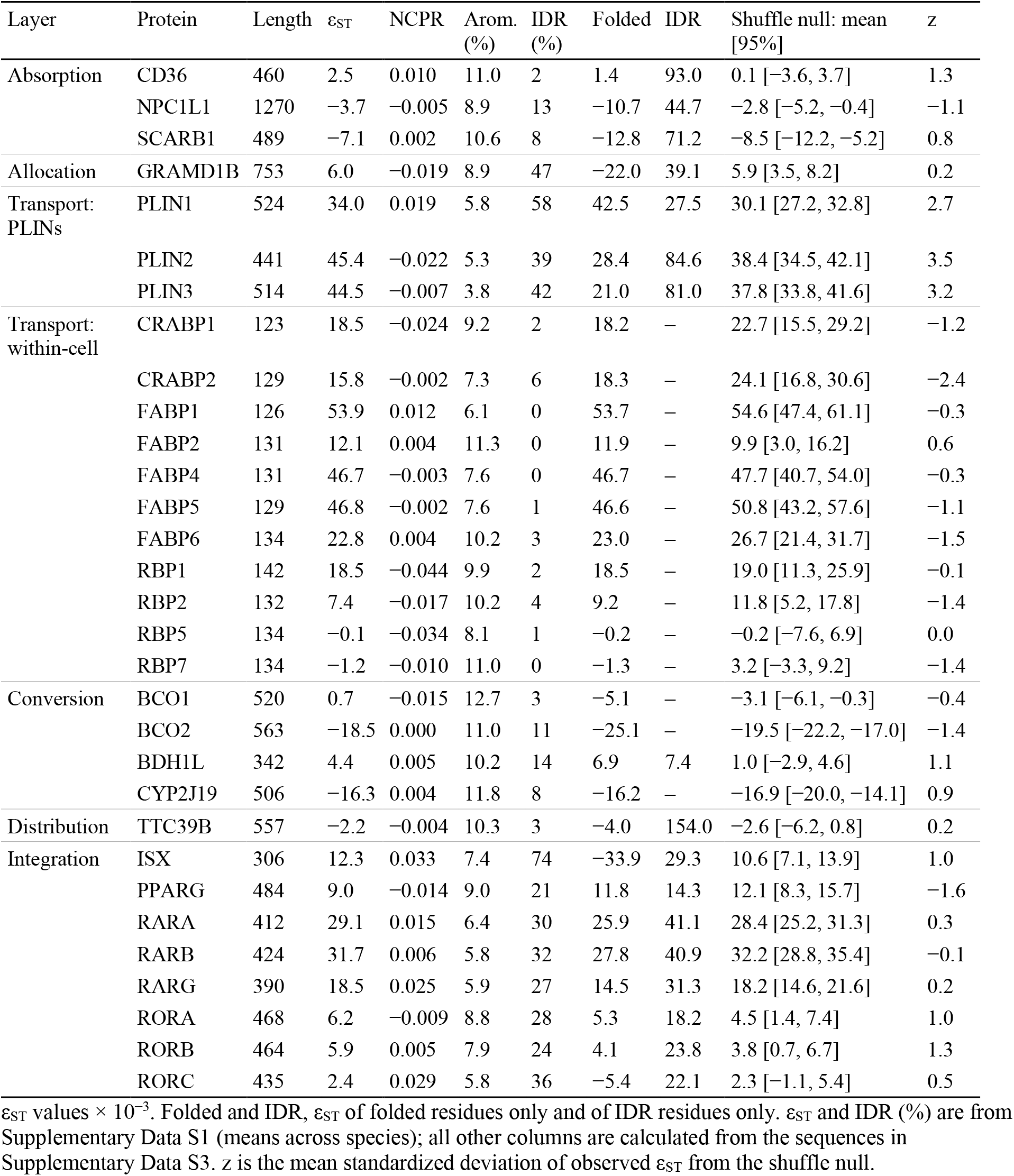
ε_ST_, sequence properties, residue subsets and shuffle null for the 31 proteins.

| Layer | Protein | Length | ε<sub>ST</sub> | NCPR | Arom. (%) | IDR (%) | Folded | IDR | Shuffle null: mean [95%] | z |
|---|---|---|---|---|---|---|---|---|---|---|
| Absorption | CD36 | 460 | 2.5 | 0.010 | 11.0 | 2 | 1.4 | 93.0 | 0.1 [−3.6, 3.7] | 1.3 |
|  | NPC1L1 | 1270 | −3.7 | −0.005 | 8.9 | 13 | −10.7 | 44.7 | −2.8 [−5.2, −0.4] | −1.1 |
|  | SCARB1 | 489 | −7.1 | 0.002 | 10.6 | 8 | −12.8 | 71.2 | −8.5 [−12.2, −5.2] | 0.8 |
| Allocation | GRAMD1B | 753 | 6.0 | −0.019 | 8.9 | 47 | −22.0 | 39.1 | 5.9 [3.5, 8.2] | 0.2 |
| Transport: PLINs | PLIN1 | 524 | 34.0 | 0.019 | 5.8 | 58 | 42.5 | 27.5 | 30.1 [27.2, 32.8] | 2.7 |
|  | PLIN2 | 441 | 45.4 | −0.022 | 5.3 | 39 | 28.4 | 84.6 | 38.4 [34.5, 42.1] | 3.5 |
|  | PLIN3 | 514 | 44.5 | −0.007 | 3.8 | 42 | 21.0 | 81.0 | 37.8 [33.8, 41.6] | 3.2 |
| Transport: within-cell | CRABP1 | 123 | 18.5 | −0.024 | 9.2 | 2 | 18.2 | – | 22.7 [15.5, 29.2] | −1.2 |
|  | CRABP2 | 129 | 15.8 | −0.002 | 7.3 | 6 | 18.3 | – | 24.1 [16.8, 30.6] | −2.4 |
|  | FABP1 | 126 | 53.9 | 0.012 | 6.1 | 0 | 53.7 | – | 54.6 [47.4, 61.1] | −0.3 |
|  | FABP2 | 131 | 12.1 | 0.004 | 11.3 | 0 | 11.9 | – | 9.9 [3.0, 16.2] | 0.6 |
|  | FABP4 | 131 | 46.7 | −0.003 | 7.6 | 0 | 46.7 | – | 47.7 [40.7, 54.0] | −0.3 |
|  | FABP5 | 129 | 46.8 | −0.002 | 7.6 | 1 | 46.6 | – | 50.8 [43.2, 57.6] | −1.1 |
|  | FABP6 | 134 | 22.8 | 0.004 | 10.2 | 3 | 23.0 | – | 26.7 [21.4, 31.7] | −1.5 |
|  | RBP1 | 142 | 18.5 | −0.044 | 9.9 | 2 | 18.5 | – | 19.0 [11.3, 25.9] | −0.1 |
|  | RBP2 | 132 | 7.4 | −0.017 | 10.2 | 4 | 9.2 | – | 11.8 [5.2, 17.8] | −1.4 |
|  | RBP5 | 134 | −0.1 | −0.034 | 8.1 | 1 | −0.2 | – | −0.2 [−7.6, 6.9] | 0.0 |
|  | RBP7 | 134 | −1.2 | −0.010 | 11.0 | 0 | −1.3 | – | 3.2 [−3.3, 9.2] | −1.4 |
| Conversion | BCO1 | 520 | 0.7 | −0.015 | 12.7 | 3 | −5.1 | – | −3.1 [−6.1, −0.3] | −0.4 |
|  | BCO2 | 563 | −18.5 | 0.000 | 11.0 | 11 | −25.1 | – | −19.5 [−22.2, −17.0] | −1.4 |
|  | BDH1L | 342 | 4.4 | 0.005 | 10.2 | 14 | 6.9 | 7.4 | 1.0 [−2.9, 4.6] | 1.1 |
|  | CYP2J19 | 506 | −16.3 | 0.004 | 11.8 | 8 | −16.2 | – | −16.9 [−20.0, −14.1] | 0.9 |
| Distribution | TTC39B | 557 | −2.2 | −0.004 | 10.3 | 3 | −4.0 | 154.0 | −2.6 [−6.2, 0.8] | 0.2 |
| Integration | ISX | 306 | 12.3 | 0.033 | 7.4 | 74 | −33.9 | 29.3 | 10.6 [7.1, 13.9] | 1.0 |
|  | PPARG | 484 | 9.0 | −0.014 | 9.0 | 21 | 11.8 | 14.3 | 12.1 [8.3, 15.7] | −1.6 |
|  | RARA | 412 | 29.1 | 0.015 | 6.4 | 30 | 25.9 | 41.1 | 28.4 [25.2, 31.3] | 0.3 |
|  | RARB | 424 | 31.7 | 0.006 | 5.8 | 32 | 27.8 | 40.9 | 32.2 [28.8, 35.4] | −0.1 |
|  | RARG | 390 | 18.5 | 0.025 | 5.9 | 27 | 14.5 | 31.3 | 18.2 [14.6, 21.6] | 0.2 |
|  | RORA | 468 | 6.2 | −0.009 | 8.8 | 28 | 5.3 | 18.2 | 4.5 [1.4, 7.4] | 1.0 |
|  | RORB | 464 | 5.9 | 0.005 | 7.9 | 24 | 4.1 | 23.8 | 3.8 [0.7, 6.7] | 1.3 |
|  | RORC | 435 | 2.4 | 0.029 | 5.8 | 36 | −5.4 | 22.1 | 2.3 [−1.1, 5.4] | 0.5 |

ε<sub>ST</sub> values × 10<sup>−3</sup>. Folded and IDR, ε<sub>ST</sub> of folded residues only and of IDR residues only. ε<sub>ST</sub> and IDR (%) are from Supplementary Data S1 (means across species); all other columns are calculated from the sequences in Supplementary Data S3. z is the mean standardized deviation of observed ε<sub>ST</sub> from the shuffle null.

**Supplementary Table S3.** Layer-level ε_ST_ for residue subsets and for the composition null.

| Layer | All residues | Shuffle null | Folded residues | IDR residues | Exposed, sequence context | Exposed, isolated beads |
|---|---|---|---|---|---|---|
| Absorption | −3.5 ± 1.2 (165) | −5.0 ± 1.3 (165) | −7.3 ± 0.9 (165) | 70.9 ± 13.6 (147) | 34.3 ± 3.0 (165) | 22.2 ± 2.9 (165) |
| Allocation | 6.4 ± 0.9 (145) | 6.2 ± 0.9 (145) | −21.9 ± 0.6 (145) | 39.3 ± 3.9 (145) | – | – |
| Transport: PLINs | 38.5 ± 1.5 (166) | 33.0 ± 1.6 (166) | 32.6 ± 1.5 (166) | 55.2 ± 4.3 (166) | – | – |
| Transport: within-cell | 14.5 ± 5.6 (82) | 17.8 ± 5.5 (82) | 14.9 ± 5.5 (82) | – | 54.9 ± 9.2 (82) | 35.7 ± 7.7 (82) |
| Transport: all carriers | 33.3 ± 1.7 (166) | 29.9 ± 1.1 (166) | 28.9 ± 1.4 (166) | 54.7 ± 4.1 (166) | – | – |
| Conversion | −12.6 ± 2.3 (31) | −13.5 ± 2.3 (31) | −11.8 ± 1.1 (31) | 7.5 ± 2.4 (16) | 25.4 ± 3.5 (14) | 5.3 ± 2.5 (14) |
| Distribution | −3.2 ± 0.8 (157) | −3.4 ± 0.7 (157) | −5.1 ± 1.0 (157) | 154.0 ± 5.6 (119) | – | – |
| Integration | 11.7 ± 0.4 (169) | 10.6 ± 0.4 (169) | 2.9 ± 2.7 (169) | 27.4 ± 5.0 (168) | – | – |

PGLS means ± s.e. × 10<sup>−3</sup> (number of species), computed from the sequences in Supplementary Data S3.

**Supplementary Table S4.** Examples of ε_ST_ of solvent-exposed and buried residues for proteins with an experimental structural template.

| Layer | Protein | Template | n | Id.<br>(%) | Exp.<br>(%) | All | Exposed<br>(context) | Exposed<br>(isolated) | Buried | RSA ≥<br>10% | RSA ≥<br>40% |
|---|---|---|---|---|---|---|---|---|---|---|---|
| Absorption | CD36 | 5LGD (H. sapiens) | 160 | 62 | 38 | 2.6 | 47.1 | 29.1 | −40.9 | −4.2 | 26.1 |
|  | SCARB1 | 5LGD (H. sapiens)* | 164 | 36 | 37 | −7.0 | 25.2 | 18.9 | −51.5 | 0.7 | 25.6 |
| Transport:<br>within-cell | CRABP1 | 2CBR (B. taurus) | 28 | 93 | 52 | 18.3 | 40.3 | 24.8 | −23.2 | 27.0 | 114.4 |
|  | CRABP2 | 1CBS (H. sapiens) | 37 | 83 | 55 | 15.5 | 31.6 | 24.8 | −27.0 | 36.7 | 52.7 |
|  | FABP1 | 3STM (H. sapiens) | 17 | 59 | 58 | 53.7 | 61.8 | 46.8 | 10.1 | 59.4 | 39.5 |
|  | FABP2 | 2IFB (R. norvegicus) | 27 | 73 | 53 | 11.9 | 69.0 | 47.8 | −59.9 | 38.4 | 110.2 |
|  | FABP4 | 3P6C (H. sapiens) | 48 | 74 | 56 | 46.7 | 86.3 | 65.8 | −29.4 | 55.1 | 60.7 |
|  | FABP5 | 1B56 (H. sapiens) | 42 | 65 | 51 | 46.8 | 84.3 | 64.9 | −13.8 | 54.4 | 76.3 |
|  | FABP6 | 1O1U (H. sapiens) | 33 | 61 | 57 | 22.7 | 63.3 | 48.7 | −63.1 | 52.0 | 51.8 |
|  | RBP1 | 1CRB (R. rattus) | 34 | 84 | 48 | 18.5 | 96.7 | 64.8 | −55.0 | 26.6 | 173.6 |
|  | RBP2 | 2RCQ (H. sapiens) | 37 | 76 | 50 | 7.4 | 63.5 | 45.9 | −49.3 | 30.3 | 37.5 |
|  | RBP5 | 1GGL (H. sapiens) | 58 | 64 | 45 | 0.0 | 19.2 | 4.9 | −19.1 | −1.9 | 105.3 |
|  | RBP7 | 1LPJ (H. sapiens) | 63 | 75 | 51 | −1.3 | 64.6 | 41.1 | −79.7 | 7.0 | 57.5 |
| Conversion | BCO1 | 3FSN (B. taurus)* | 14 | 43 | 29 | −3.7 | 50.5 | 21.4 | −37.2 | 0.5 | 22.4 |
|  | BCO2 | 3FSN (B. taurus)* | 12 | 41 | 31 | −21.4 | −8.6 | −16.6 | −30.3 | −22.4 | −15.5 |

ε<sub>ST</sub> values × 10<sup>−3</sup>, means across species; n, number of species. Templates: PDB entry (source organism); asterisks mark paralogous templates (CD36 for SCARB1; RPE65 for BCO1 and BCO2). Id., mean sequence identity between avian sequences and the template over aligned positions. Exp., percentage of aligned residues with RSA ≥ 25%. All, all residues; Exposed, residues with RSA ≥ 25% with sequence-context weights or as isolated beads; Buried, residues with RSA < 25%. The last two columns give exposed residues at alternative RSA thresholds (isolated beads).

